# Proteomic profiling of primary cilia in the developing brain uncovers new regulators of cortical development

**DOI:** 10.1101/2025.05.03.652041

**Authors:** Xiaoliang Liu, Oscar Torres Gutierrez, Gurleen Kaur, Yazan Al-Issa, Sabyasachi Baboo, Jolene K. Diedrich, Eva Cai, John R. Yates, Xuecai Ge

## Abstract

In the developing brain, the primary cilia of radial glial cells extend from the surface of the lateral ventricle, serving as the signaling hub to integrate environmental cues critical for brain formation. Dysfunctions in ciliary proteins contribute to a wide range of brain structural abnormalities in ciliopathies, yet identifying the bona fide ciliary protein components in the brain remains a significant challenge. Here, using proximity labeling and quantitative proteomics, we systematically charted proteins localized to the cilia of radial glial cells in the dorsal and ventral regions of the embryonic brain. From this dataset, we identified cohorts of new molecules intrinsic to the cilia at distinct regions of the developing brain. We validated the cilium localization of several components of the translation machinery and studied the mechanistic roles of ciliary candidates previously linked to brain malformation, including Marcks, a key regulator of radial glial polarity, and Ckap2l, a protein associated with Filippi syndrome. These results revealed previously unrecognized ciliary mechanisms that regulate brain development. Thus, our brain proteomic dataset provides a unique resource for understanding ciliary functions in brain development and the molecular etiology of developmental disorders.

## INTRODUCTION

The primary cilium is a microtubule-based sensory organelle projecting from the cell surface. Often referred to as the cell’s antenna, the primary cilium senses a diverse array of signaling cues from the surrounding environment and transduces them into intracellular signaling pathways that elicit specific cellular responses^1^. Unique receptors and signaling transducers are concentrated in the primary cilia of different organs, enabling cells to respond to specific environmental cues^2,3^. Dysfunction of primary cilia leads to a wide range of diseases, collectively known as ciliopathies^4^. Brain structural abnormalities, such as microcephaly, neuronal heterotopia, disrupted cortical lamination, and cerebellar hypoplasia, are common features of ciliopathies^5–10^. These structural deficits typically arise from abnormal brain development during embryogenesis^11^. However, the precise roles of primary cilia in cortical formation remain underexplored.

The embryonic neurogenesis of the mammalian brain begins with radial glia (RG), which produce most neurons in the cortex. Over the course of embryonic neurogenesis, RG strike an intricate balance between self-renewal and neuronal production^12^. They first proliferate to expand the neural progenitor pool, followed by several rounds of neurogenic divisions to produce neurons. At the end of neurogenesis, the RG population is depleted due to neurogenic division and differentiation into astrocytes, oligodendrocytes or ependymal cells^13,14^. How RG achieve this intricate balance is not fully understood. An intriguing yet under-investigated mechanism is that RG are subject to regulatory molecular cues from cerebrospinal fluid (CSF) that circulates signaling molecules in the developing central nervous system^15^. The primary cilia of RG extend from the ventricular surface and are immersed in CSF, ideally positioned to detect the signaling molecules^15–17^. A systematic study on the *bona fide* protein components in the cilia of RG is promising to provide critical insights, but such a study has yet to be conducted.

RG in the developing brain constitute a highly heterogeneous population, displaying significant diversity across and within different brain regions^18–22^. In the dorsal and ventral cortex, RG exhibit distinct molecular profiles and contribute differently to development. Dorsal RG, which express markers such as PAX6 and SOX2, are primarily responsible for generating excitatory projection neurons^23^. These neurons migrate radially to the cortical plate, forming cortical layers in an inside-out manner^24^. In contrast, ventral RG, located in the lateral and medial ganglionic eminences (LGE and MGE), give rise to inhibitory neurons^25,26^. These inhibitory neurons migrate tangentially into the dorsal cortex, where they integrate into circuits with the excitatory projection neurons^27^. In addition, within the same brain regions, RG exhibit considerable heterogeneity. In the dorsal cortex, subpopulations of radial glia display variations in gene expression, proliferative capacity, and differentiation potential^28^, revealed by recent single-cell RNA sequencing studies^29,30^. This heterogeneity of RG is essential for the precise regulation of neurogenesis over space and time, enabling the establishment of complex neural circuits. Such differential cell fate determination is typically governed by a combination of extrinsic signaling cues and intrinsic transcriptional programs^31–33^. However, the specific external cell signaling that contributes to the heterogeneity of RG remains poorly understood.

Primary cilium functions as a signaling hub for the cell, prompting us to determine the authentic ciliary proteins in RG. A comprehensive map of ciliary proteins in RG could provide crucial insights into the extrinsic signaling mechanisms that regulate RG specification and their role in brain development. This is a challenging task because the primary cilium constitutes ∼1/10,000 of the total cell volume; conventional approaches of physically isolating cilia usually contain contaminations from non-ciliary structures. In addition, some ciliary proteins are present in the cilium at very low abundance, and most signaling transducers transit through the cilia in a very dynamic manner. Physical isolation is not able to capture this dynamic protein presence in the cilium. Proximity labeling strategies have proven effective to identify ciliary proteins in cultured cells^34,35^. In a previous study, we employed TurboID, an engineered biotin ligase, to profile new ciliary proteins involved in Hedgehog (Hh) signaling^36^. This approach allowed us to capture low-abundant proteins in the cilia in response to Hh pathway activation.

Here, we leveraged TurboID-based proximity labeling to systematically map ciliary proteins in the RG of the mouse embryonic brain. We generated a Cilium-TurboID transgenic mouse model and performed quantitative ciliary proteomics in over 800 embryonic brains, providing both broad and deep coverage of ciliary proteome during brain development. Our results revealed region-specific differences in ciliary composition between the dorsal and ventral cortex. Further, our dataset uncovered molecular machinery for local protein synthesis in the cilia of RG, suggesting an unexpected layer of ciliary regulation. Finally, our proteomic analysis identified a set of ciliary proteins linked to neurodevelopmental disorders, leading to the discovery of previously unrecognized ciliary mechanisms underlying these conditions. Together, our dataset provides a unique resource for exploring the understudied biology of primary cilia in the developing brain and the molecular etiology of neurodevelopmental disorders.

## RESULTS

### Constructing a mouse model for *in vivo* biotinylation of ciliary proteins in RG in the developing brain

To selectively label proteins in the primary cilia of RG, we generated a transgenic mouse line in which the transgene Cilium-TurboID is specifically targeted to the cilia, and its expression is driven by the BLBP promoter (Figure 1A). The transgenic mice were generated via the piggyBac transposon system^37^. In the transgene, TurboID is fused to a truncated form of ARL13B (ι1ARL13B), which was created in our previous study and has been characterized to specifically localize to the cilium without disturbing ciliary morphology and function^36^. The BLBP promotor is active in RG from the early developmental stage (E10.5) throughout cortical development^38,39^.

**Figure 1.**
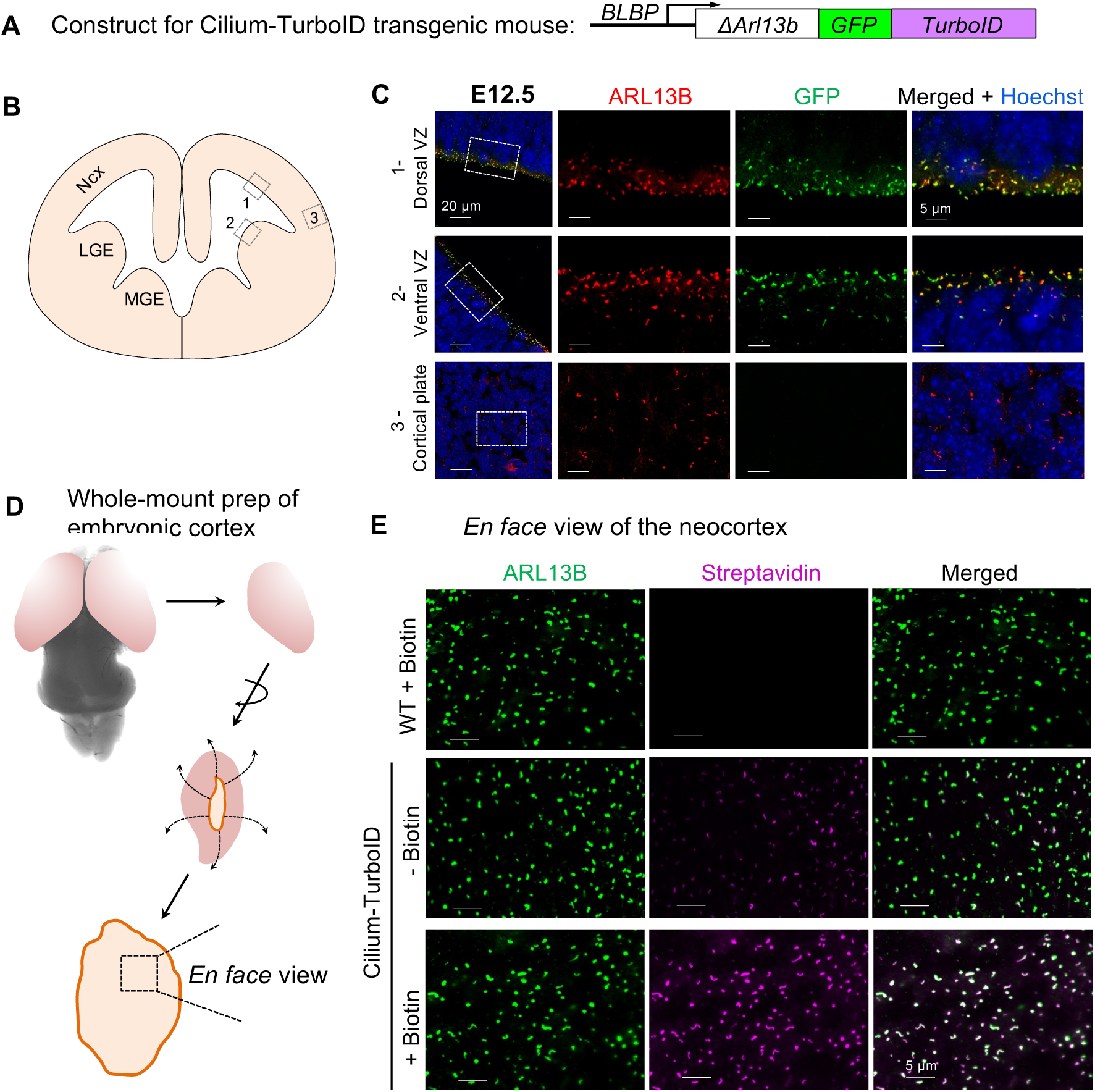
Constructing a mouse model for *in vivo* biotinylation of ciliary proteins in RG in the developing brain. (**A**) The schematic of the transgenic construct used for the generation of Cilium-TurboID mouse model. The transgene is expressed as one fused protein of Λ1ARL13B-GFP-TurboID. Λ1ARL13B is a truncated form of ARL13B, containing only the N+RVEP+PR domain of ARL13B. (**B**) The schematic of the embryonic brain section. Ncx, neocortex; LGE, lateral ganglionic eminence; MGE, medial ganglionic eminence. (**C**) Immunofluorescence of cortical sections from dorsal and ventral brain regions of E12.5 Cilium-TurboID transgenic embryos. The transgene was detected by immunostaining for GFP (green); cilia are labelled by ARL13B (red); DNA is visualized by Hoechst staining (blue). Areas highlighted by gray dashed boxes in different regions are magnified and displayed at the right side. (**D**) Diagram illustrating the whole-mount preparation of the embryonic cortex. The two hemispheres are separated, and the hemispheres are cut radially around the ventral opening. The inner surface of the cortex was imaged directly from an *en face* perspective. (**E**) Whole-mount staining of transgenic mouse neocortex expressing Cilium-TurboID or WT mouse neocortex before and after biotin labeling. E12.5 brains were incubated with 50 µM biotin for 15 min at 37°C, and fixed for immunostaining. The cilium is labeled by ARL13B (green) and the biotinylated proteins were labeled by streptavidin-Alexa Fluor 647 (magenta). Scale bars, 5 µm in all panels in (**C**) and (**E**).

The transgene has no discernible effect on the development or function of any organs, and the brain size and morphology are indistinguishable from those of WT mice (data not shown). In this transgenic mouse line, we first confirmed that Cilium-TurboID was expressed in RG within the embryonic brain. Immunofluorescence staining on coronal sections shows that the transgene is specifically expressed in all primary cilia of RG at the ventricular surface across both dorsal and ventral regions, but not in the cilia of mature neurons in the cortical plate (Figure 1B and 1C). To validate the activity of TurboID in the cilium, we incubated the freshly dissected embryonic cortices in biotin, and prepared whole-mount brain tissues by exposing the inner surface of the cortex. We then directly imaged the primary cilia from the *en face* perspective (Figure 1D). We found that all primary cilia at the ventricular surface of the transgenic mouse brain were strongly labeled with biotin after a 15-minute incubation with biotin (Figure 1E). We detected a slight background biotinylation signal in the cilia of the transgenic brain in the absence of biotin incubation. In contrast, no biotin signal was detected in the cilia of wild-type (WT) brain.

To enrich biotinylated proteins from the mouse brain, we tested beads coated with either streptavidin or neutravidin. Both proteins bind biotin with extremely high affinity (Kd ≈ 10^-14^ to 10^-15^ M); however, compared to streptavidin, neutravidin exhibits lower non-specific binding due to its neutral charge, and the absence of glycosylation and the RYD sequence ^40–42^. Indeed, we found that the overall protein intensities enriched by streptavidin beads are indistinguishable between WT and Cilium-TurboID brains. However, with the same amount of total protein input, neutravidin beads enriched higher protein intensities from the Cilium-TurboID brains compared to the WT brains (Figure S1A and S1B). We hence used neutravidin beads to enrich biotinylated proteins in the subsequent proteomic assays.

### *In vivo* ciliary proteomics in the developing mouse brain

Using the Cilium-TurboID transgenic mouse model, we sought to chart the profile of ciliary proteins in RG with quantitative proteomics in two brain regions: the dorsal region (neocortex) and the ventral region (ganglionic eminence) (Figure 2A). We focused on embryonic day 12.5 (E12.5), a critical stage when the RG population is abundant and actively balancing neurogenesis with self-amplification. For each labeling condition (WT + biotin, Transgenic + biotin, Transgenic + vehicle), 420-430 embryonic brains were collected. An equal amount of 14 mg total protein input per condition was used, evenly divided into three replicates for enrichment with neutravidin beads (Figure S2A). The enriched proteins were digested on the beads and tagged with a 10plex Tandem Mass Tag (TMT) label set. The multiplexed samples were then analyzed using HPLC-MS/MS for protein identification and quantification (Figure 2A).

**Figure 2.**
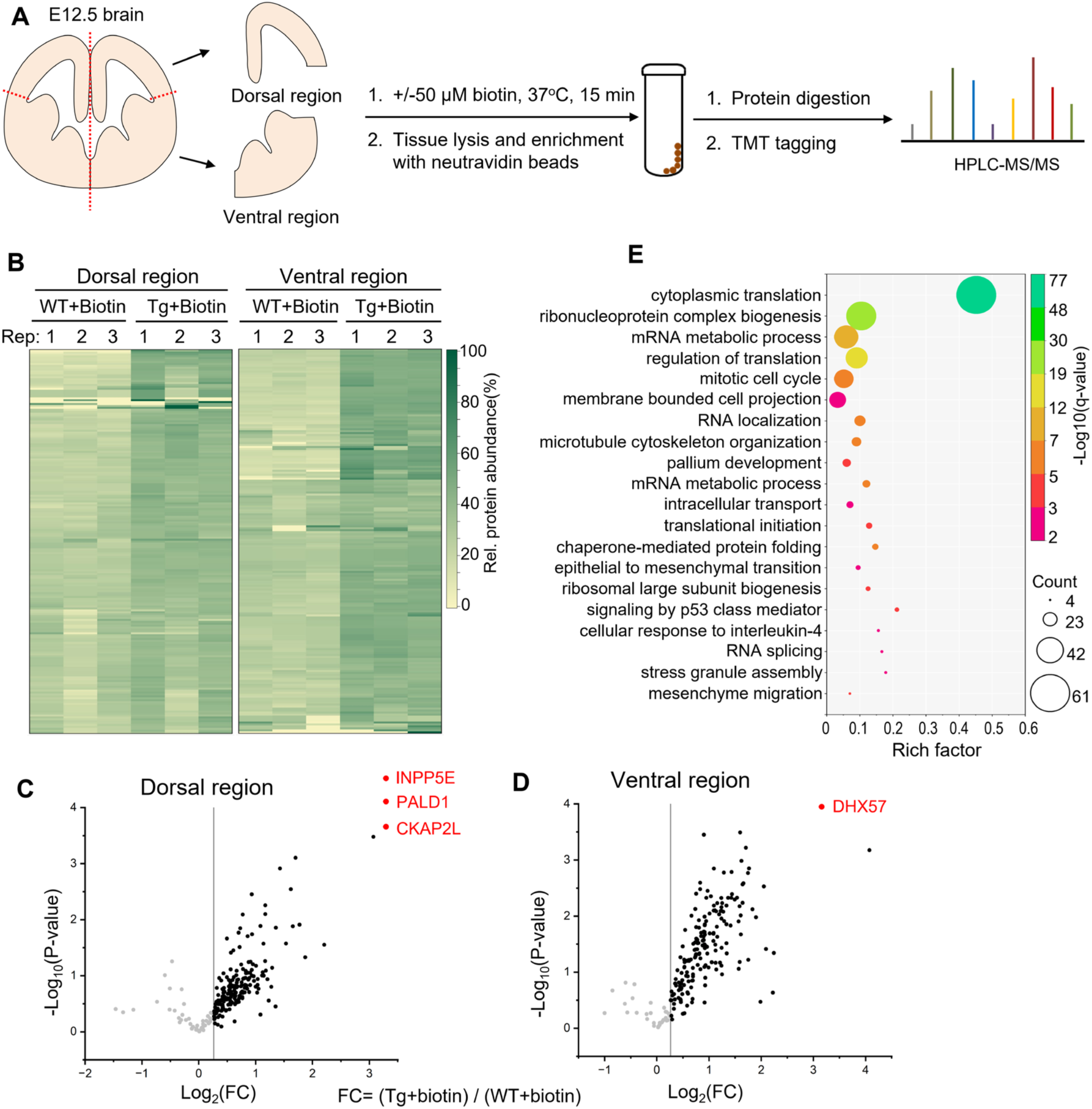
*In vivo* ciliary proteomics of the developing mouse brain. (**A**) Workflow of in vivo ciliary proteomic study in the embryonic mouse brains. For each brain, one hemisphere was used for biotin labeling, and the other for vehicle control. The dorsal and ventral cortex are separated and processed in parallel. (**B**) Heatmap of the relative abundance of ciliary candidates in the dorsal and ventral brain region. Clustering of the relative abundances of each identified protein (rows) in individual samples (columns) was performed based on Ward’s minimum variance method. The relative abundance was calculated within one set of samples that contain all three conditions. The panel on the right side indicates the color scheme of relative abundances (percentage). (**C**) Volcano plot depicting statistical significance versus the average enrichment ratio of ciliary candidate proteins from the dorsal brain region. The enrichment ratio was determined by comparing the TMT signal intensity of proteins in the biotin-labeled transgenic brain to those in the WT brain. Candidate proteins highlighted in red (INPP5E, PALD1, and CKAP2L) were exclusively detected in the transgenic brain and were absent in the WT brain. (**D**) Volcano plot depicting statistical significance versus the average enrichment ratio of ciliary candidate proteins from the ventral brain region. The candidate highlighted in red (DHX57) was exclusively detected in the transgenic brain and was absent in the WT brain. (**E**) GO enrichment analysis of biological processes was plotted according to the rich factor in Metascape. The top 20 enriched biological processes are represented in the bubble plot.

For data analysis, we defined the relative protein abundance as the ratio of normalized abundance in each channel over the reference channel. The heatmap of each protein’s relative abundance and the correlation of biological replicates demonstrated high reproducibility across the triplicates (Figure 2B and S2B). We then took the ratio of protein abundance in the TMT channel of the biotin-labeled transgenic brain over that in the biotin-labeled WT brain. The volcano plots were generated depicting statistical significance versus the averaged enrichment ratio (Figure 2C-D). With a cut-off of TMT ratio > 1.2, we obtained 198 and 191 ciliary candidate proteins from the dorsal and ventral brain regions, respectively. By combining these two sets, we identified a total of 258 proteins as ciliary candidates, independent of their region of origin. Among them, 33 candidates have been previously reported to associate with cilia, including ciliary signaling regulators (INPP5E, PALD1, GSK3B, IRS1), proteins involved in ciliary structure and ciliogenesis (Tubulin proteins, CFAP20, FLNA, FLNB), and these associated with protein sorting and trafficking (TULP3, UNC119, PACS1, KIF5B, RAN) (Figure S2C).

Gene Ontology (GO) analysis of Biological Processes, along with KEGG and CORUM pathway analysis, reveals that the 258 ciliary candidates participate in diverse pathways and biological processes (Figure 2E). In addition to expected terms such as cytoskeleton organization, intracellular transport, and pallium development, the top 20 enriched clusters include unexpected categories, such as protein translation regulation, RNA binding, and RNA localization. In the following studies, we sought to validate this dataset by confirming the ciliary localization of unexpected ciliary candidates and determining the ciliary functions of proteins previously linked to brain developmental disorders.

### Embryonic brain ciliary proteomics uncover translation machinery components in the cilia of radial glia

Out of the 258 ciliary candidates in our dataset, 68 proteins (26%) belong to ribosome components and translation factors (Figure S3A). Notably, these proteins were also uncovered as ciliary candidates in three previous proteomic studies in other systems, such as IMCD3 cells^43^, marine models^44^, and mouse ependymal cells^45^ (Figure 3A). Further, the study on ependymal cells demonstrated that local protein synthesis supports the structure and function of motile cilia^45^. Yet, mRNA localization and local protein synthesis have not been identified in primary cilia. This prompted us to validate the presence of translational machinery and mRNA in the primary cilium.

**Figure 3.**
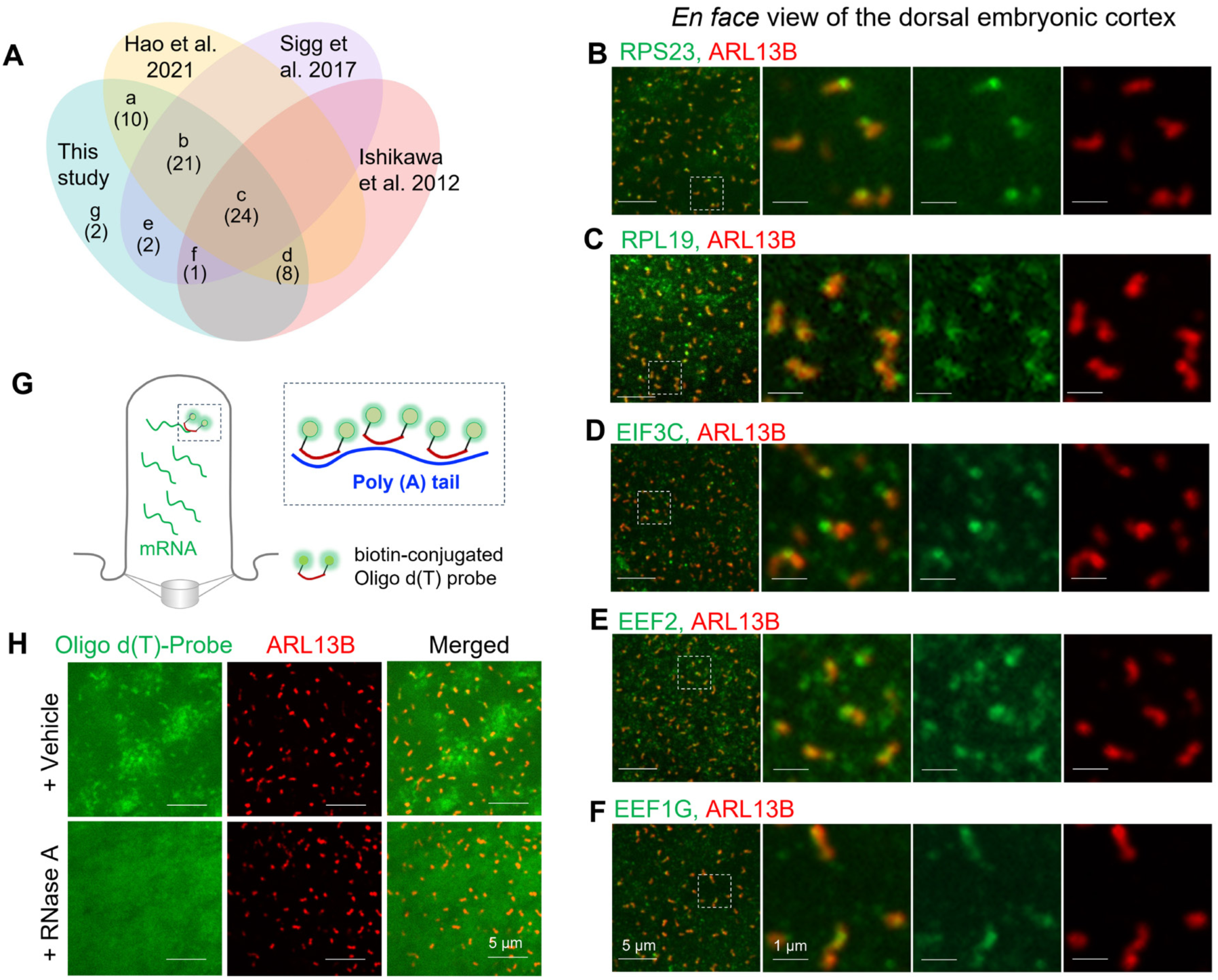
Embryonic brain ciliary proteomics uncover translation machinery components in the cilia of radial glia. (**A**) *Venn* diagram showing overlapping of translation machinery components identified in three previous ciliary proteomic studies and our current study. The list of overlapping proteins is included in Table S2. (**B**-**F**) *En face* view of the dorsal cortex in the whole-mount E12.5 mouse brain. The brain tissues were immuno-stained with antibodies against the indicated translation machinery proteins. Primary cilia are labeled with ARL13B (red). Regions within the white dashed boxes are magnified and shown on the right side. (**G**) Diagram illustrating fluorescence in situ hybridization (FISH) of ciliary mRNA using poly-d (T) probes conjugated with biotin. The signaling was detected via tyramide signal amplification (TSA). (**H**) Representative images of FISH on whole-mount E12.5 mouse cortices. Cilia were imaged from an *en face* perspective. Immunostaining with ARL13B was performed to highlight the cilium (red).

We performed immunofluorescence staining on whole-mount mouse cortical preparation and imaged primary cilia from the *en face* perspective (Figure S3B). The results demonstrate that multiple components of translation machinery are present in the cilia of RG, including ribosome components, RPS23 and RPL19, and translation factors, EIF3C, EEF2 and EEF1G (Figure 3B-F and S3C-F). Certain ribosome components, such as RPL23A, specifically localize to the base of cilia in RG (Figure S3G-H). Other ribosome components, such as RPS3, RPL11, and RPL10A have previously been identified in the motile cilia of brain ependymal cells^45^. However, in RG cells, their high signal levels in the cell body precluded us from resolving individual cilia in the whole-mount cortices. Nevertheless, we validated their presence in the cilia of NIH3T3 cells (Figure S3I-K).

Next, we sought to detect the presence of mRNA in the cilia of RG. On the whole-mount mouse cortices, we conducted RNA fluorescence *in situ* hybridization (FISH) using a biotin-conjugated Oligo(dT) probe, which was subsequently detected with streptavidin-HRP followed by an HRP substrate reaction (Figure 3G). Simultaneously, we carried out immunofluorescence staining to label the cilia. Our results revealed a robust Oligo(dT) signal in the cilia of RG, suggesting the presence of polyadenylated mRNA (Figure 3H). Further, this signal was abolished by RNase A treatment, confirming its RNA specificity.

Taken together, our results demonstrate the presence of mRNA and translation machinery in the cilia of RG in the embryonic cortex, suggesting a potential role of local protein synthesis in the developing brain.

### Identifying new ciliary function for neurodevelopmental disorder-associated proteins

Ciliary dysfunction is widely implicated in neurodevelopmental disorders (NDDs)^6,46^. To explore this connection, we compared our brain ciliary proteomic dataset with curated gene sets associated with NDDs, including Prebirth BrainSpan, SynGO, and GeneTrek^47^. 204 ciliary candidates in our dataset overlap with at least one of these resources, and 46 candidates overlap with all three (Figure 4A). To gain further insights, we focused on the candidates in the 46 proteins that have well-established functions in brain development but have not yet been characterized as ciliary proteins. Among them, MARCKS (Myristoylated alanine-rich C-kinase substrate) emerges as a top candidate.

**Figure 4.**
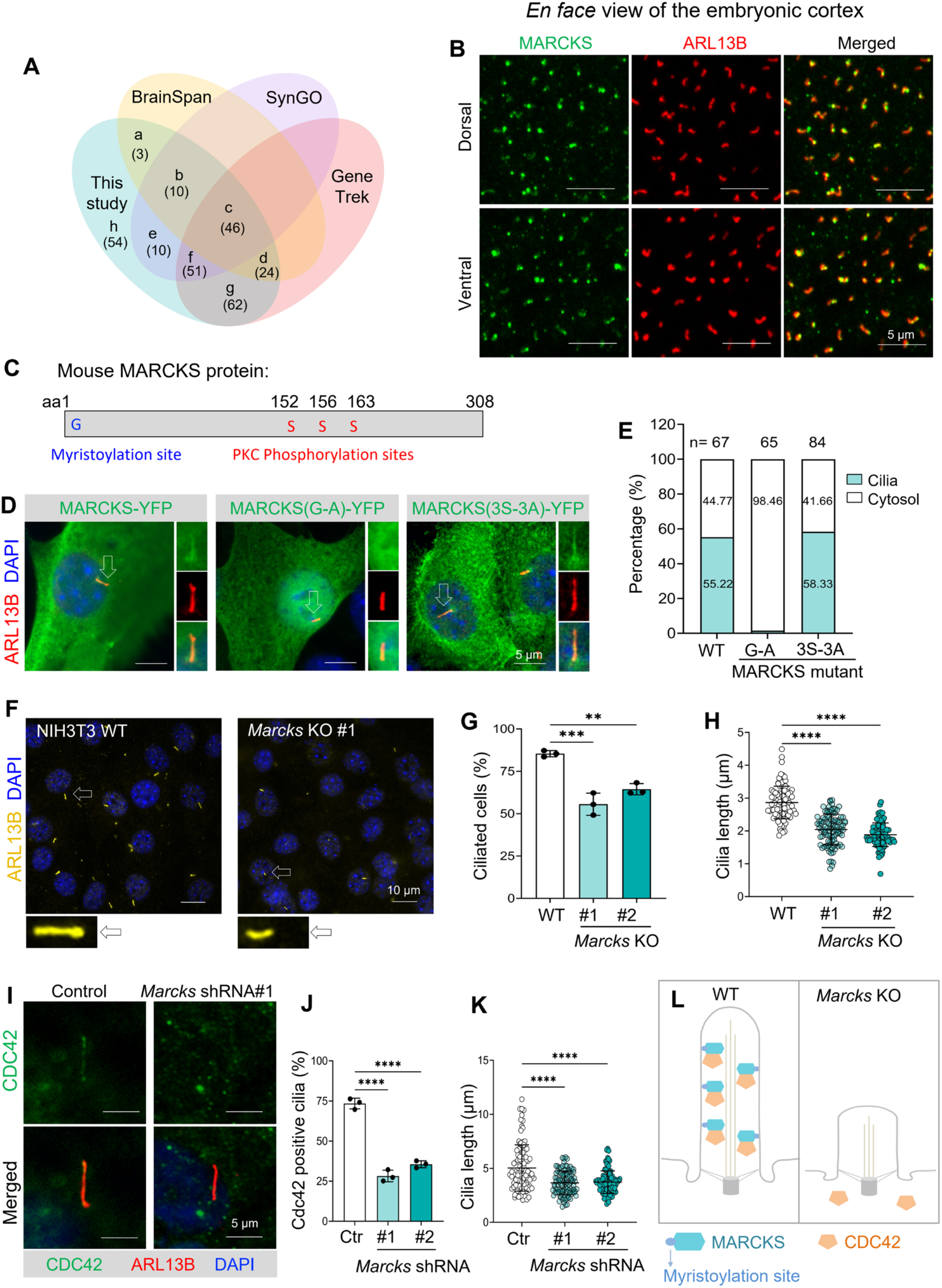
*In vivo* ciliary proteomics Identified new ciliary function for neurodevelopmental disorder-associated proteins. (**A**) *Venn* diagram showing common proteins shared in our brain ciliary proteomic datasets and other three curated databases. The list of overlapping proteins is included in Table S3. (**B**) Representative images showing that endogenous MARCKS proteins are present in the primary cilia of E12.5 mouse cortex in both the dorsal and ventral regions. (**C**) A schematic of full-length MARCKS protein, with the myristoylation site and PKC phosphorylation sites highlighted. (**D**) Myristoylation is required for the cilium localization of MARCKS in NIH3T3 cells. YFP-tagged WT or mutated MARCKS were expressed in NIH3T3, and their subcellular localization were determined by immunofluorescence staining. The cilium pointed by the white arrow is magnified and displayed at the right side. (**E**) Quantification of cilium localization for WT and mutant MARCKS. The total number of cells quantified for each condition was listed in the histogram. Experiment was performed 3 times with similar results. (**F**) Representative images of cilium staining with ARL13B antibody in WT or *Marcks* KO cells. The cilium pointed by the white arrow is magnified and displayed at the bottom. (**G**-**H**) Quantification of ciliation rate (**G**) and cilium length (**F**) in WT or *Marcks* KO cells. For each condition, at least 100 cilia were quantified. Experiments were performed 3 times with similar results. Data are presented as mean ± SD. Statistics: One-way ANOVA with multiple comparisons (Tukey test). **, p < 0.01; ***, p < 0.001; ****, p < 0.0001. (**I**) Representative images showing ciliary CDC42 levels in control and *Marcks* knockdown ARPE-19 cells. Cells were infected with lentiviruses expressing shRNA targeting *Marcks*. Three days post-infection, cells were serum-starved for 48 hours to induce ciliogenesis, followed by immunofluorescence staining. (**J**) Quantification of cilia that are positive for CDC42 immunofluorescence signal in WT or *Marcks* knockdown ARPE-19 cells. (**K**) Quantification of ciliary length in WT or *Marcks* knockdown ARPE-19 cells. In (**J**-**K**), at least 100 cilia were quantified for each experimental condition. Experiment was performed 3 times with similar results. Data are presented as mean ± SD. Statistics: One-way ANOVA with multiple comparisons (Tukey test). ****, p < 0.0001. (**L**) Schematic of role of MARCKS in targeting CDC42 during ciliogenesis and ciliary growth. MARCKS anchors to the ciliary membrane via myristoylation site, which then recruits CDC42. The ciliary targeted CDC42 is essential for maintaining the vesicle trafficking required for ciliogenesis and ciliary extension. In the absence of MARCKS, cells either fail to form a cilium or exhibit shortened cilia.

MARCKS is a membrane-associated protein and a putative substrate of protein kinase C (PKC)^48,49^. Loss of *Marcks* leads to severe neurodevelopmental defects, including failed neural tube closure, incomplete brain hemisphere separation, and disorganization of the RG scaffold^50–53^. The underlying molecular mechanisms remain poorly understood. To identify the role of MARCKS in the primary cilia, we first validated its ciliary localization. MARCKS-YFP localizes to the cilium when expressed in NIH3T3 cells. Further, immunostaining with a MARCKS antibody revealed its presence in the cilia in various cell types, including NIH 3T3, SH-SY5Y, and human RPE cells (Figure S4A). Immunostaining on whole-mount embryonic cortices shows that MARCKS is present in the primary cilia of RG across dorsal and ventral regions (Figure 4B).

Previous studies have shown that the membrane association of MARCKS is mediated by the myristoylation of its first glycine residue (G1)^54–56^. In addition, PKC phosphorylation has been mapped to three serine residues (S152,S156, S163) in its internal domain^57^ (Figure 4C). Genetic studies showed that the myristoylation modification, but not PKC phosphorylation, is needed for MARCKS’ role in radial glial scaffold organization^52^. To determine whether these post-translational modifications regulate the ciliary localization of MARCKS, we constructed mutants that abolish either myristoylation (G-A mutation) or phosphorylation (3S-3A mutation) and assessed their subcellular localization in NIH3T3 cells. We found that the myristoylation mutant completely lost its localization to the primary cilia, whereas the phosphorylation mutant retained ciliary localization similar to WT MARCKS (Figure 4D-E). Hence, MARCKS’ ciliary localization depends on its myristoylation but does not require PKC phosphorylation.

To identify the functional role of MARCKS in the primary cilium, we generated *Marcks* knockout cells via CRISPR/Cas9 in NIH3T3 cells. Western blot results confirmed its depletion at the protein level (Figure S4C-D). *Marcks* knockout significantly reduced the ciliation rate. Moreover, in the remaining ciliated cells, the ciliary length was dramatically shortened compared to WT cells (Figure 4F-H). To unravel the underlying mechanism of MARCKS in ciliogenesis, we focused on its previously reported role in targeting cell polarity-related proteins, such as CDC42^52^. CDC42 facilitates vesicular trafficking essential for delivering components to the growing cilium^58–60^, and its levels in the cilium serve as an indicator of its activity during this process. To test it, we silenced *Marcks* expression with shRNA in a human RPE cell line, ARPE-19 (Figure S4E), and assessed CDC42 levels in the cilium. We found that *Marcks* knockdown significantly reduced CDC42 levels in the cilium (Figure 4I-J). The cilium length in RPE cells is also significantly shortened by *Marcks* knockdown (Figure 4K).

Collectively, these data suggest that loss of *Marcks* blocks the targeting of CDC42 to ciliogenesis sites, thereby impairing the vesicular transport essential for cilium assembly. As a result, cells exhibit either absent or shortened cilia (Figure 4L). It is likely that the neurodevelopmental defects observed in *Marcks* knockout mice are partially linked to defective ciliogenesis and disrupted ciliary function in neural progenitors during brain development.

### Brain cilium proteomics uncover region-specific variations in ciliary composition of RG

Our cilium proteomic dataset indicates an intriguing difference in the protein compositions of cilia between the dorsal and ventral brain regions (Figure 2B-D). Further analysis identified 67 proteins specifically enriched in the dorsal region, 60 in the ventral region, and 131 proteins common to both regions (Figure 5A).

**Figure 5.**
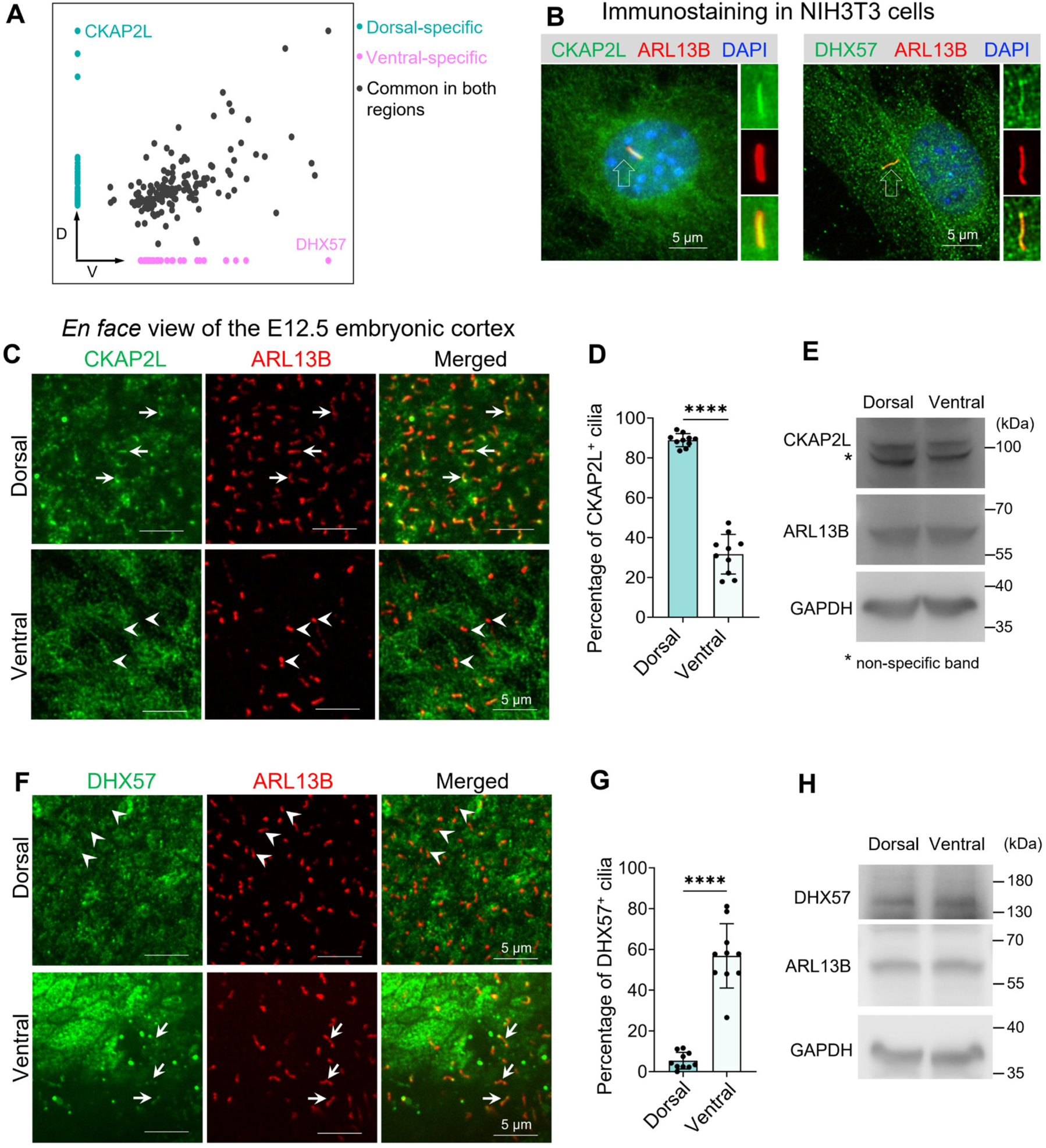
Brain cilium proteomics uncover region-specific variations in ciliary composition of RG. (**A**) Scatter plot of ciliary candidates in dorsal versus ventral brain regions. The dorsal-specific, ventral-specific, and common ciliary candidates in both regions are listed in Table S4. (**B**) Representative images of immunofluorescence staining showing that CKAP2L and DHX57 are present in the primary cilia of NIH3T3 cells. The cilium pointed by the white arrow is magnified and displayed at the right side. (**C**) *En face* views of whole-mount embryonic mouse brain following immunofluorescence staining with an anti-CKAP2L antibody. White arrows indicate representative cilia positive for CKAP2L in the dorsal region; white arrowheads denote cilia lacking CKAP2L immunoreactivity in the ventral region. (**D**) Quantification of CKAP2L-positive cilia in (**C**). 10 areas in each brain regions from three mice were quantified. (**E**) Western blotting results of CKAP2L expression levels in the dorsal and ventral brain regions of E12.5 mouse. (**F**) *En face* views of whole-mount embryonic mouse brain following immunofluorescence staining with an anti-DHX57 antibody. White arrows indicate representative cilia positive for DHX57 in the ventral region; white arrowheads denote cilia lacking DHX57 immunoreactivity in the dorsal region. (**G**) Quantification of DHX57-positive cilia in (**F**). 10 areas in each brain regions from three mice were quantified. (**H**) Western blotting results of DHX57 expression levels in the dorsal and ventral brain regions of E12.5 mouse. Data in (**D**) and (**G**) are presented as mean ± SD. Statistics: two-tailed Student’s t-test. ****, p < 0.0001.

To further investigate region-specific ciliary localization, we validated the top candidates: Cytoskeleton Associated Protein 2-Like (CKAP2L) in the dorsal cortex, and DExH-Box Helicase 57 (DHX57) in the ventral cortex. Notably, cilium localization of these two proteins has not been previously reported. Immunostaining in NIH3T3 cells shows that both proteins are present in the cilium (Figure 5B). To determine their localization in the cilia of RG, we prepared immunofluorescence staining on whole-mount E12.5 cortices. We mounted the dorsal and ventral (LGE and MGE) cortex separately based on their relative positions and surface contours (Figure S3B). We then imaged the cilia of RG at the inner ventricular surface from the *en face* perspective. We found that CKAP2L is present in the majority of cilia in the dorsal cortex (88.9 ± 3.3%), whereas only a small portion of cilia in the ventral cortex contain CKAP2L (31.6 ± 9.39%) (Figure 5C-D). In contrast, DHX57 is present in most cilia in the ventral cortex (56.9 ± 15.9%) and only a small fraction of cilia in the dorsal cortex (5.4 ± 4.1%) (Figure 5F-G). Intriguingly, Western blot analysis reveals that the total expression levels of both proteins are comparable between the dorsal and ventral cortex (Figure 5E and 5H).

These results suggest that the region-specific ciliary localization is not due to variations in overall protein expression, but rather reflects distinct regulatory mechanisms that govern protein targeting to cilia in different brain regions. In addition, the reciprocal ciliary localization patterns of CKAP2L and DHX57 validate our initial findings in the proteomic dataset, highlighting region-specific variations in ciliary composition between the dorsal and ventral brain regions.

### CKAP2L regulates cilium-dependent cell signaling

Next, we characterized roles of new ciliary candidates from our proteomic dataset in cilium-dependent cell signaling, focusing on CKAP2L. Loss-of-function mutations in CKAP2L lead to Filippi Syndrome, a rare genetic disorder characterized by short stature, microcephaly, intellectual disability, and syndactyly^61–66^. CKAP2L was originally identified as a mitotic spindle-associated protein in neural progenitors of both the embryonic and adult brain^67^. In addition to mitotic spindles in dividing cells (data not shown), we found that CKAP2L localizes to the primary cilia of interphase cells in multiple cell types, including radial glia (Figure 5C and S5A), NIH3T3 cells (Figure 5B), SH-SY5Y and human RPE cells (Figure S5A). Expansion microscopy reveals that CKAP2L distributes along the ciliary axoneme, consistent with its known role as a microtubule-associated protein^68^ (Figure S5B).

To determine the mechanistic roles of CKAP2L, we generated *Ckap2l* knockout (KO) NIH3T3 cells using CRISPR/Cas9 (Figure S5D). Western blot analysis with a validated CKAP2L antibody confirmed efficient depletion of CKAP2L proteins (Figure 6A and S5E). *Ckap2l* KO cells exhibit no abnormalities in cilium length or ciliation rate, and the chromosome segregation and cell cycle progression remain normal (data not shown). Next, we determined its role in ciliary signal transduction. The Hh pathway is the most well-characterized cilium-dependent signaling, and the symptoms of Filippi syndrome patients resemble defects in Hh signaling^61,69^. Therefore, we assessed Hh signaling activity in *Ckap2l* KO cells after stimulation with the Sonic Hedgehog (Shh) ligand. qPCR analysis on two Hh pathway target genes, *Gli1* and *Ptch1*, showed that the Shh-induced transcript levels of both genes were markedly reduced in *Ckap2l* KO cells compared to those in WT cells (Figure 6B-C). These results suggest attenuated Hh signaling in *Ckap2l* KO cells.

**Figure 6.**
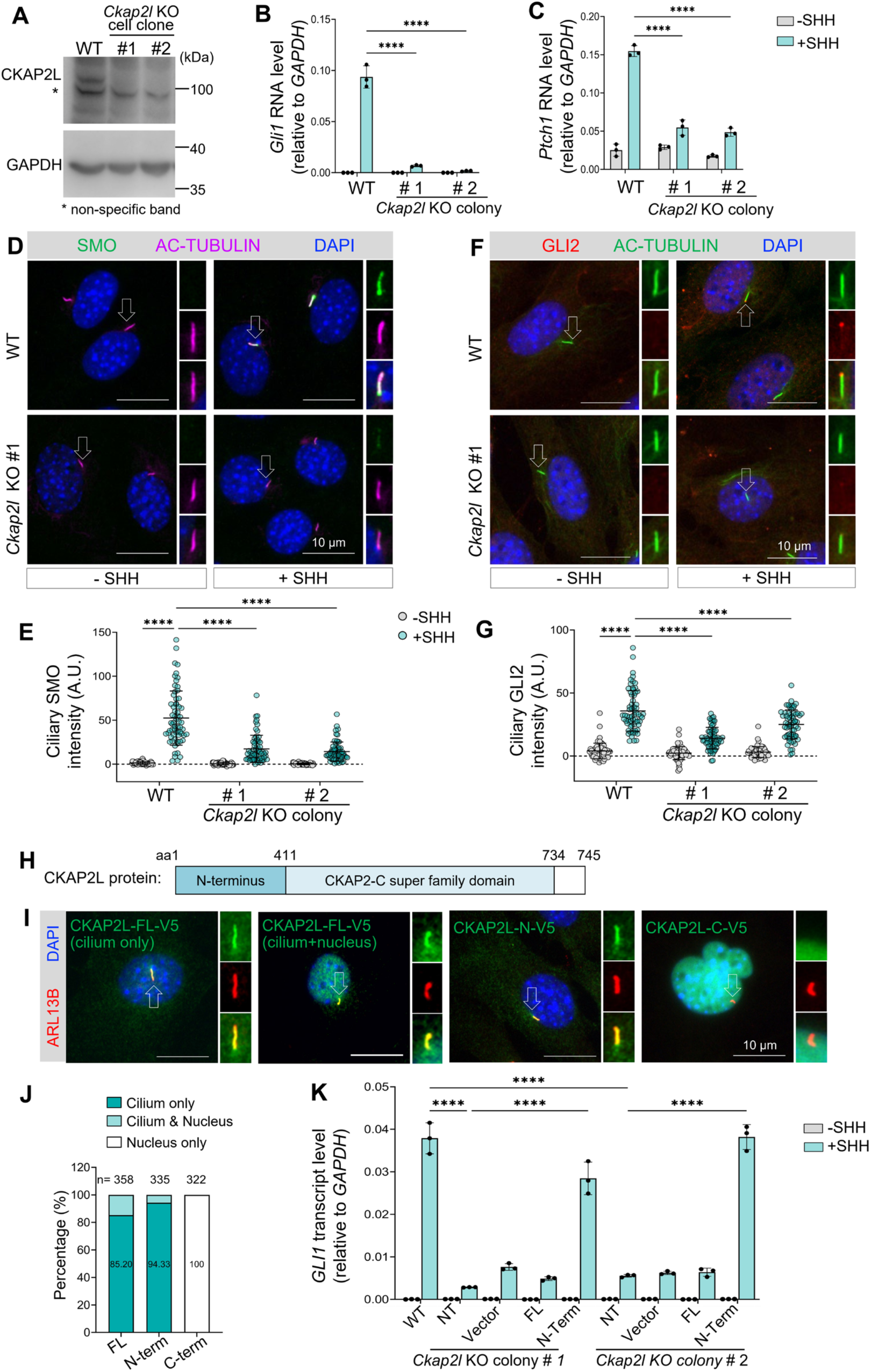
CKAP2L regulates cilium-dependent cell signaling. (**A**) Western blot analysis showing the absence of CKAP2L protein in the *Ckap2l* CRISPR/Cas9 knockout NIH3T3 cell clones. Asterisk indicates a non-specific band. (**B**-**C**) Hh signaling levels in WT or *Ckap2l* KO NIH3T3 cells were assessed by the transcript levels of the Hh-target genes, *Gli1* and *Ptch1*, via qPCR. Data are shown as mean ± SD. Statistics: Two-way ANOVA with multiple comparisons (Tukey test), ****, p < 0.0001. (**D**) Immunofluorescence of SMO in the cilia in WT and *Ckap2l* KO NIH3T3 cells. Cells were treated with vehicle or SHH for 24 h before being fixed for staining. The cilium pointed by the white arrow is magnified and displayed at the right side. (**E**) Quantification of SMO fluorescence intensity in the cilium. A total of 65 cilia were quantified for each experimental condition. A.U., Arbitrary Unit. Data are shown as mean ± SD. Statistics: Two-way ANOVA with multiple comparisons (Tukey test), ****, p < 0.0001. (**F**) Immunofluorescence staining of endogenous GLI2 in the cilia of WT or *Ckap2l* KO NIH3T3 cells. Cells were treated with vehicle or SHH for 24 h before being fixed for staining. The cilium pointed by the white arrow is magnified and displayed at the right side. (**G**) Quantification of GLI2 fluorescence intensity at the cilium tips. A total of 65 cilia were quantified for each experimental condition. A.U., Arbitrary Unit. Data are shown as mean ± SD. Statistics: Two-way ANOVA with multiple comparisons (Tukey test), ****, p < 0.0001. (**H**) Diagram showing the CKAP2L protein structure. (**I**) Representative images showing localization of CKAP2L full length (FL), N- and C-terminus in NIH3T3 cells. (**J**) Percentage of cells expressing the indicated construct localized to the cilium or nucleus. (**K**) Hh signaling levels in WT cells and *Ckap2l* KO NIH3T3 cells that express full-length CKAP2L, N-terminal form or the blank vector. Hh activity was assessed by *Gli1* transcript levels. Data are shown as mean ± SD. Statistics: Two-way ANOVA with multiple comparisons (Tukey test), ****, p < 0.0001.

The Hh signaling is transduced across the membrane by Smoothened (SMO), a GPCR-like receptor that is translocated and activated in the cilium. Active SMO then triggers a signaling cascade that eventually leads to the activation of transcription factor GLI2 at the ciliary tip^70^. To determine whether CKAP2L is required for this process, we assessed SMO and GLI2 ciliary levels in *Ckap2l* KO cells via immunofluorescence staining. We found that Shh-induced ciliary accumulation of both SMO and GLI2 is significantly reduced in *Ckap2l* KO cells compared to WT cells (Figure 6D-G). Thus, CKAP2L positively regulates Hh signaling by facilitating SMO translocation to the cilium, which subsequently promotes GLI2 activation.

To exclude potential off-target effects of CRISPR/Cas9, we re-introduced CKAP2L protein in *Ckap2l* KO cells via lentivirus-mediated gene expression. Unfortunately, CKAP2L-full length leads to disorganized mitotic spindles and distorted nuclei in a significant percentage of cells, as reported in a previous study^67^ (Figure S5C), preventing the proper assessment of Hh signal activity. The CKAP2L protein features a CKAP2-super family domain at the C-terminus and a more diversified N-terminus (Figure 6H). Interestingly, we found that the N-terminal domain predominantly localizes to the primary cilium, whereas the C-terminal domain is exclusively present in the nucleus (Figure 6I-J). Further, expressing the C-terminal domain led to disorganized nuclei in nearly all infected cells. Finally, expressing the N-terminal domain in *Ckap2l* KO cells was sufficient to restore Shh-induced Hh signaling (Figure 6K). These results indicate that CKAP2L’s ciliary and nuclear functions are separatable: the N-terminus alone suffices to regulate Hh signaling in the cilium, while the C-terminus acts in the nucleus.

### *Ckap2l* loss disrupts brain development and Hh signaling in the developing cortex

To determine the roles of CKAP2L in brain development, we generated genetic a mouse model of *Ckap2l* knockout via CRISPR/Cas9 (Figure S6A-B). Western blot analysis on the cortices confirmed complete the removal of CKAP2L from the homozygous brain (Figure S6C). The *Ckap2l* KO mice (*Ckap2l^-/-^*) are born at the expected Mendelian ratio. However, they appear runty, exhibit reduced body size, and most are unable to breed. The brain size in adult *Ckap2l* KO mice is reduced across multiple dimensions (Figure S7A-F). Sagittal sections of the brain at the same mediolateral level show that most brain regions are smaller in *Ckap2l* KO mice compared to WT mice, with pronounced underdevelopment in the cortex and most cerebellar lobules (Figure S7G).

As *Ckap2l* loss in NIH3T3 cells leads to reduced Hh signaling (Figure 6), we tested Hh signaling levels in the developing cortex. Among the three GLI transcription factors, GLI3 plays a predominant role in dorsal telencephalon development and is essential for balancing neural progenitor proliferation and differentiation^71–73^. Hh signaling is suppressed by GLI3R, a proteolytic product of GLI3 full length. Activation of the Hh pathway ceases the GLI3 proteolytic cleavage, reducing GLI3R production. The relative abundance of GLI3R often serves as an indicator of Hh signaling intensity. We therefore examined the GLI3R levels in the dorsal and ventral regions of WT and *Ckap2l^-/-^* brains via Western blot. We found a significant increase in Gli3R levels in the dorsal region of *Ckap2l^-/-^* mice compared to the WT mice (Figure S7H-I). However, the GLI3R levels in the ventral regions were comparable between WT and *kap2l^-/-^* brains. This result aligns with diminished Hh signaling observed in *Ckap2l* KO NIH3T3 cells. Further, the dorsal-specific effect on Hh signaling corroborates our findings that CKAP2L is primarily localized to cilia in the dorsal but not the ventral cortex (Figure 5).

### *Ckap2l* loss reduces embryonic neurogenesis in the developing brain

In the developing telencephalon, Hh signaling is essential for maintaining the neural progenitor pool in the dorsal cortex, and its reduction leads to decreased cortical size^74,75^. Therefore, we examined neural progenitor populations in the developing cortex using immunostaining for SOX2, a marker of radial glia, and TBR2, a marker of intermediate neural progenitors (INPs). In E13.5 *Ckap2l^-/-^* mice, both SOX2-positive and TBR2-positive layers were significantly thinner compared to WT mice. In addition, the overall thickness of the cerebral cortex was reduced in *Ckap2l^-/-^* embryos relative to WT controls (Figure 7A-B). Thus, *Ckap2l* loss reduces the neural progenitor pool in the developing dorsal cortex.

**Figure 7.**
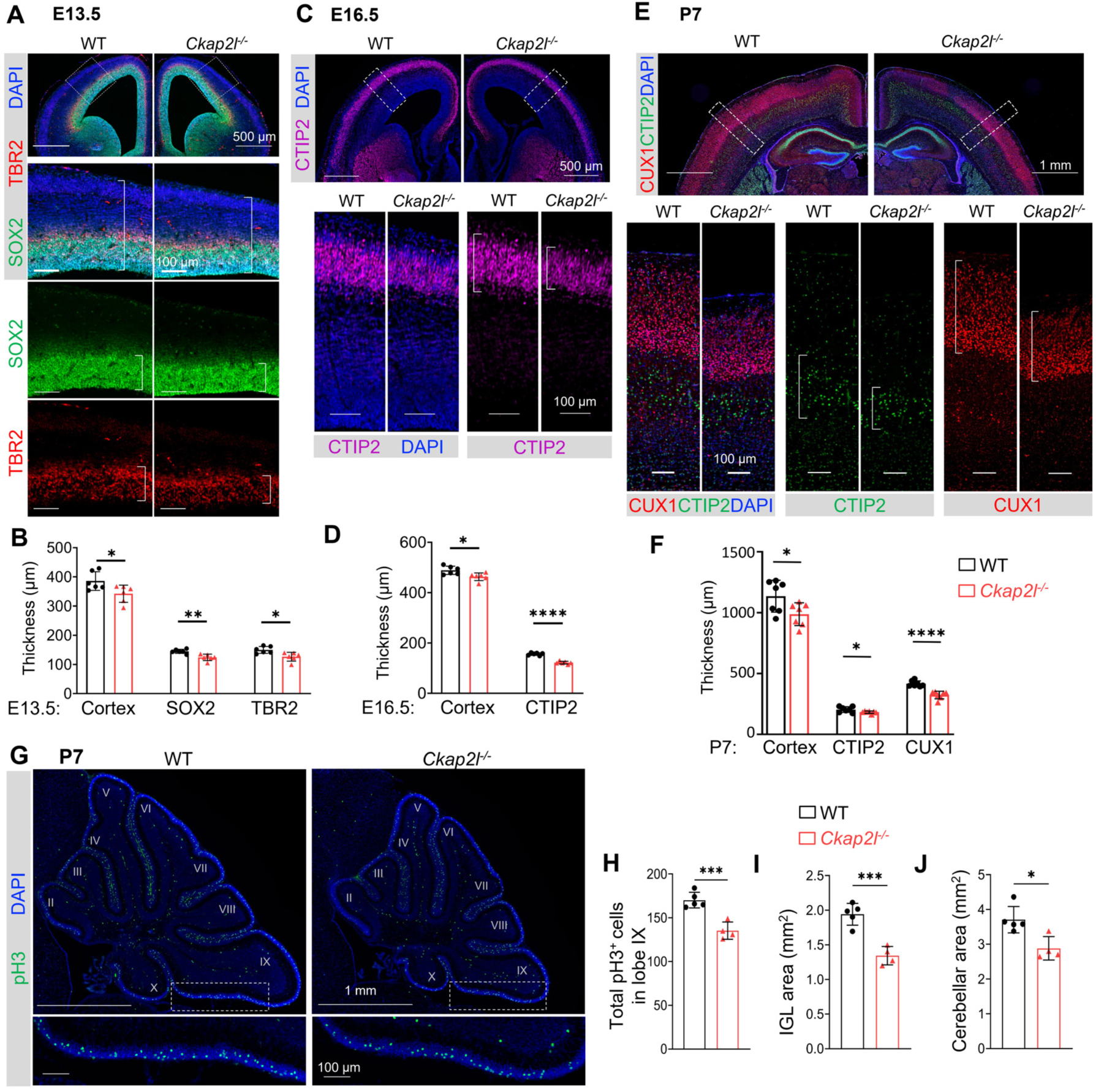
*Ckap2l^-/-^* mouse exhibits reduced neurogenesis. (**A**) Representative images of E13.5 brain sections stained with SOX2 and TBR2. Areas in white dashed boxes are magnified and displayed at the bottom. White brackets demarcate the thickness of the indicated layer. (**B**) Quantification of the thickness of indicated layers in (**A**). Coronal sections at the same rostral-caudal level indicated in (**A**) were used for quantification. (**C**) Representative images of E16.5 brain sections stained with CTIP2. Areas in white dashed boxes are magnified and displayed at the bottom. (**D**) Quantification of the thickness of indicated layers in (**C**). Coronal sections at the same rostral-caudal level indicated in (**C**) were used for quantification. (**E**) Representative images of P7 brain sections stained with CTIP2 and CUX1. Areas in white dashed boxes are magnified and displayed at the bottom. (**F**) Quantification of the thickness of indicated layers in (**E**). Coronal sections at the same rostral-caudal level indicated in (**E**) were used for quantification. For (**B**), (**D**), and (**F**), n=6-7 brains were quantified for each genotype. Data are presented as mean ± SD. Statistics: two-tailed Student’s t-test. ****, p < 0.0001; **, p < 0.01; *, p < 0.05. (**G**) Sagittal sections of cerebellum at the same medio-lateral levels from P7 WT and *Ckap2l^-/-^* mice. Sections were immune-stained with anti-pH3 antibody. White dashed boxes are magnified and displayed at the bottom. Lobes are indicated by Roman number. (**H**) Quantification of the number of pH3-positive cells in the EGL in lobe IX in the sections of the same medio-lateral levels. (**I**-**J**) Quantification of the IGL area and cerebellar area in P7 mice measured at the same medio-lateral level. For (**H**-**J**), n = 4 or 5 cerebella were quantified for each genotype. Data are presented as mean ± SD. Statistics: two-tailed Student’s t-test. ***, p < 0.001; *, p < 0.05.

Next, we determined the impact of *Ckap2l* loss on neurogenesis. By E16.5, when early born CTIP2-positive neurons are generated, we found a reduced population of CTIP2-positive neurons in *Ckap2l^-/-^* embryos compared to WT littermates (Figure 6C-D). This reduction is not due to developmental delay, as the CTIP2-positive neuronal layer remains thinner in *Ckap2l^-/-^* cortices at postnatal stages (P3 and P7). Furthermore, the layer of later-born CUX1-positive neurons is also thinner in the *Ckap2l^-/-^* brain compared to the WT littermates (Figure 7E-F and S8A-B). Finally, this reduction in neuronal populations is consistently observed across the entire rostral-caudal axis (Figure S8C-D). Overall, these results suggest that *Ckap2l* loss reduced neural progenitor pools, leading to diminished neuronal production across all developmental stages, which ultimately results in reduced cortical size.

Shh functions as a critical mitogen for granule neuron precursors (GNPs) in the developing cerebellum^76–78^. Given that CKAP2L positively regulates Hh signaling, we next assessed GNP proliferation in the developing cerebellum. Immunostaining with the mitotic marker phospho-histone H3 (pH3) revealed a reduced number of proliferating cells in the external granule layer (EGL) of *Ckap2l^-/-^* cerebella compared to WT littermates (Figure 7G-H). Differentiated granule neurons migrate from the EGL to the internal granule layer (IGL); we further examined cerebellar architecture and found that both the IGL area and the total cerebellar area were significantly reduced in *Ckap2l^-/-^* cerebella compared to the WT littermates (Figure 7I-J). Thus, loss of *Ckap2l* reduced GNP proliferation, leading to cerebellar hypoplasia.

While the *Ckap2l* KO mouse model exhibits key features of Filippi syndrome such as short stature and microcephaly, it fails to fully recapitulate other symptoms, such as syndactyly. Potential functional redundancy could be provided by other similar proteins, such as CKAP2, a paralog of CKAP2L. The two proteins share 16% amino acid identity and a C-terminal CKAP2 super-family domain^79,80^. We found that CKAP2 localizes to the primary cilia of multiple cell types, including NIH3T3, SH-SY5Y, and radial glial cells (Figure S9A-B). Whole-mount cortical staining further confirmed CKAP2’s presence in the cilia of both the dorsal and ventral cortex (Figure S9C-D). To elucidate CKAP2’s function, we used shRNA-mediated knockdown in NIH3T3 cells. While CKAP2 depletion did not affect ciliogenesis or ciliary length (Figure S9E-G), it significantly attenuated Shh-induced Hh pathway activation, as indicated by reduced *Gli1* transcript level (Figure S9H). Thus, CKAP2 acts as a positive regulator of Hh signaling and may functionally compensate for CKAP2L loss in certain contexts.

## DISCUSSION

Brain structural deficits are common in ciliopathies, underscoring the critical roles of primary cilia in brain development^5–10^. Primary cilia are present in various cell types in the developing brain, including migrating newborn neurons, mature neurons, and neural progenitors. Ciliary defects likely disrupt key biological processes across all these cell types. In this study, we focused on cilia in RG and defined their *bona fide* ciliary protein composition using TurboID-mediated proximity labeling and quantitative proteomics. A number of previous studies using proximity labeling enzymes, such as BioID/BioID2, APEX/APEX2, and TurboID, have markedly expanded the repertoire of ciliary proteins^34,81–83^. However, it is important to note that these studies were conducted in cultured cell lines. The *in vitro* systems lack the multicellular interactions between diverse cell types and the dynamic signaling cues, such as morphogen gradients and fluid flow, that shape ciliary function and protein composition *in vivo*. Therefore, *in vivo* studies are essential for accurately capturing the physiological complexity of the ciliary proteome in brain development. Indeed, beyond known ciliary proteins (Figure S2C), we identified a set of novel ciliary candidates and uncovered the mechanistic roles of several in ciliary signaling and brain development. Our study represents the first comprehensive dataset of brain ciliary proteomics, and establishes a valuable resource for understanding cilium-mediated cell signaling in RG and the molecular basis of ciliopathies and brain developmental disorders.

Intriguingly, our study revealed region-specific variation in the ciliary composition of RG between the dorsal and ventral cortex. We validated this regional specificity by confirming CKAP2L ciliary localization in the dorsal cortex and DHX57 in the ventral cortex (Figure 5). While primary cilia across different cell types share core structural components, their functional protein composition can vary significantly depending on cell types and contexts. For instance, PDGFRA localizes to cilia in fibroblasts but not in RG^36,84,85^. Similarly, neuronal cilia may express unique receptors, such as NPY2R, which is absent in epithelial cells^86,87^. And SMO is detectable in the cilium only upon activation of Hh signaling^87,88^. Many factors could contribute to the differential ciliary localization, such as cell-type-specific gene expression, post-translational modifications, variations in ciliary trafficking machinery, and differential exposure to external signaling pathways. It is well-established that RG in the dorsal and ventral cortex represent distinct cell populations^18–22^ and they may also be exposed to different external signaling cues. Understanding the mechanisms that drive differential ciliary targeting in distinct RG subtypes and its functional implications highlights important directions for future study.

Our *in vivo* ciliary proteomics identified a set of translation machinery components, including ribosomal subunits and eukaryotic initiation and elongation factors (EIFs and EEFs) (Figures 3 and S3). While many of these components have been detected in previous ciliary proteomes, none have been validated in the primary cilia^36,43,44^. Here, we provide direct evidence for the presence of translation machinery, as well as mRNAs, in the primary cilia (Figure 3). These findings strongly support the presence of local protein synthesis in primary cilia. This concept appears to challenge the prevailing view that the intraflagellar transport (IFT) machinery is primarily responsible for delivering proteins to the cilium for assembly and maintenance^89^. Nevertheless, several notions suggest that local translation within cilia could be necessary under specific conditions.

First, local translation has been demonstrated in the motile cilia of brain ependymal cells, where it is essential for maintaining the proper ciliary function^45^. This finding supports the feasibility of protein synthesis within the ciliary microenvironment. Second, one of the translation factors validated in our study, EIF3, has been reported to directly regulate Hh signaling. Loss-of-function mutations in *Eif3* cause the *extra-toes spotting* (*Xs*) in mice, which is characterized by polydactyly—a hallmark of excessive Hh pathway activation^90^. Intriguingly, *Xs* mutants show no global translation defects but exhibit specific protein reduction in PTCH1, a key negative regulator of Hh signaling^90,91^. Further study revealed that EIF3C binds to a pyrimidine-rich motif, which is present in the 5’ UTRs of *PTCH1* transcripts^91^. Together, the evidence suggests that EIF3C may regulate the local translation of Hh pathway components in the cilium, thereby modulating Hh signaling on-site. Finally, local translation is best established in neuronal growth cones and synapses, structures that are distal to the cell body and require on-site protein synthesis for rapid, spatially restricted responses^92–95^. Notably, RG exhibit a similar spatial constraint: their primary cilia remain anchored at the apical surface while the nucleus undergoes interkinetic nuclear migration (INM), oscillating between the apical and basal boundary of the ventricular zone in coordination with the cell cycle^96,97^. As a result, the cilium and nucleus are often physically separated. Local translation in the cilia of RG may enable direct synthesis of structural or signaling proteins to ensure timely responses to extracellular cues. For future studies, it will be intriguing to identify ciliary mRNAs in RG and determine how localized translation contributes to cortical development and function.

Our *in vivo* ciliary proteomics offered new insights into the molecular underpinnings of neurodevelopmental disorders. For example, we uncovered a ciliary role for MARCKS, a protein whose critical functions in embryonic mouse brain development were identified decades ago, yet its molecular mechanisms remain poorly understood^50–53^. Although MARCKS is a well-defined PKC substrate with conserved phosphorylation sites, genetic studies have shown that these sites are dispensable for brain development. Instead, its N-terminal myristoylation is essential^52,54–57^. Intriguingly, we found that the myristoylation site, but not the PKC phosphorylation sites, is required for MARCKS ciliary localization. These findings suggest that the ciliary function of MARCKS is fundamental during developing brain. Further, MARCKS plays a key role in ciliogenesis by recruiting CDC42 and facilitating CDC42-associated vesicle transport to support cilium formation (Figure 4F-L). Thus, the neurodevelopmental defects in *Marcks* mutants may result from impaired ciliogenesis and subsequent disruptions of ciliary signaling.

CKAP2L is another previously uncharacterized ciliary protein implicated in neurodevelopmental disorders. Loss-of-function mutations in CKAP2L lead to Filippi syndrome. We thoroughly characterized the cilium localization of CKAP2L, and pinpointed its N-terminus as the ciliary targeting domain and the previously described nuclear localization domain to the C-terminus. Consistent with previous studies, we found that over-expression of CKAP2L, particularly the C-terminus, disrupted nuclear morphology by interfering with mitotic spindle assembly and chromosome segregation (Figure 6I and S5C) ^67^. Unlike previous reports, we found that CKAP2L loss-of-functions does not disrupt chromosome segregation, instead, it impaired Hh signal transduction by restricting the ciliary translocation of SMO and subsequent GLI2 activation (Figure 6D-G). Importantly, CKAP2L is present in nearly all cilia in the dorsal cortex but only a subset in the ventral cortex. While the underlying mechanism requires future inquiry, this cell-type specific cilium localization may explain the moderate phenotype in *Ckap2l* KO mice. Hh signaling functions as a morphogen that specifies ventral cell fates in the neural tube, and its reduction leads to dorsalization^98^. Surprisingly, *Ckap2l* KO mice show no obvious neural patterning defects, likely because CKAP2L is largely absent from ventral cilia, sparing Hh signaling in this region. However, after neural patterning, dorsal Hh signaling is critical to maintain RG and INP neural progenitor pools^74,75^. Consequently, *Ckap2l* loss disrupts neuronal populations in both the outer and inner cortical plates. Notably, *Ckap2l* KO mice do not recapitulate syndactyly observed in human Filippi syndrome patients. While such mouse-human discrepancies are common, functional compensation by its paralog, CKAP2, may explain this difference. Like CKAP2L, CKAP2 localizes to primary cilia and promotes Hh signaling (Figure S9), potentially buffering the loss of CKAP2L in mice.

Our in vivo ciliary proteomics study offers a unique dataset of the *bona fide* ciliary proteome in the developing mammalian brain. We identified previously uncharacterized ciliary proteins, expanding our understanding of the molecular landscape of primary cilia in neurodevelopment. This dataset serves as a valuable resource for the broader scientific community. Future studies leveraging this resource could further elucidate cilium-mediated signaling mechanisms and their roles in neurodevelopmental pathologies.

## KEY RESOURCES TABLE

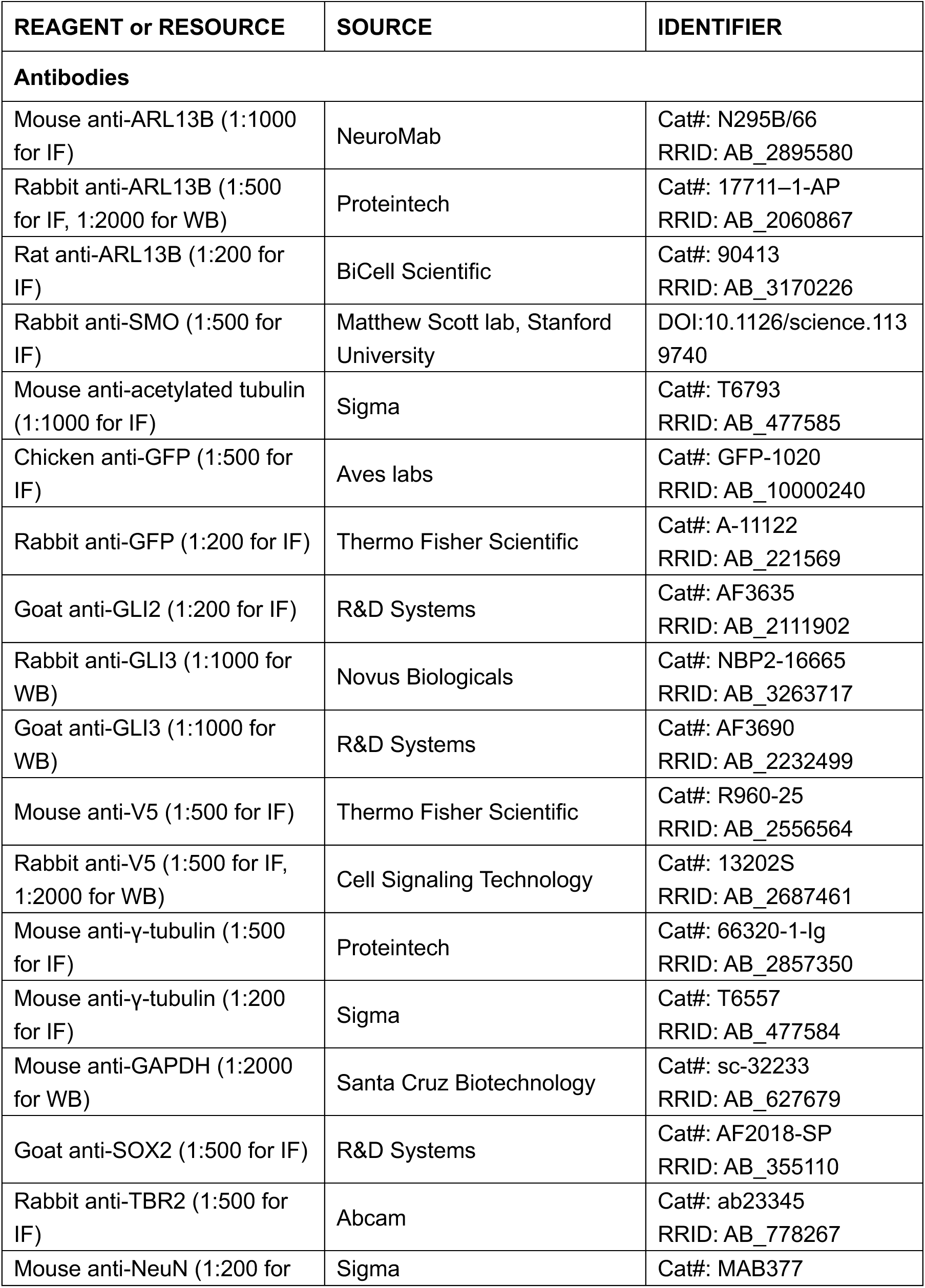
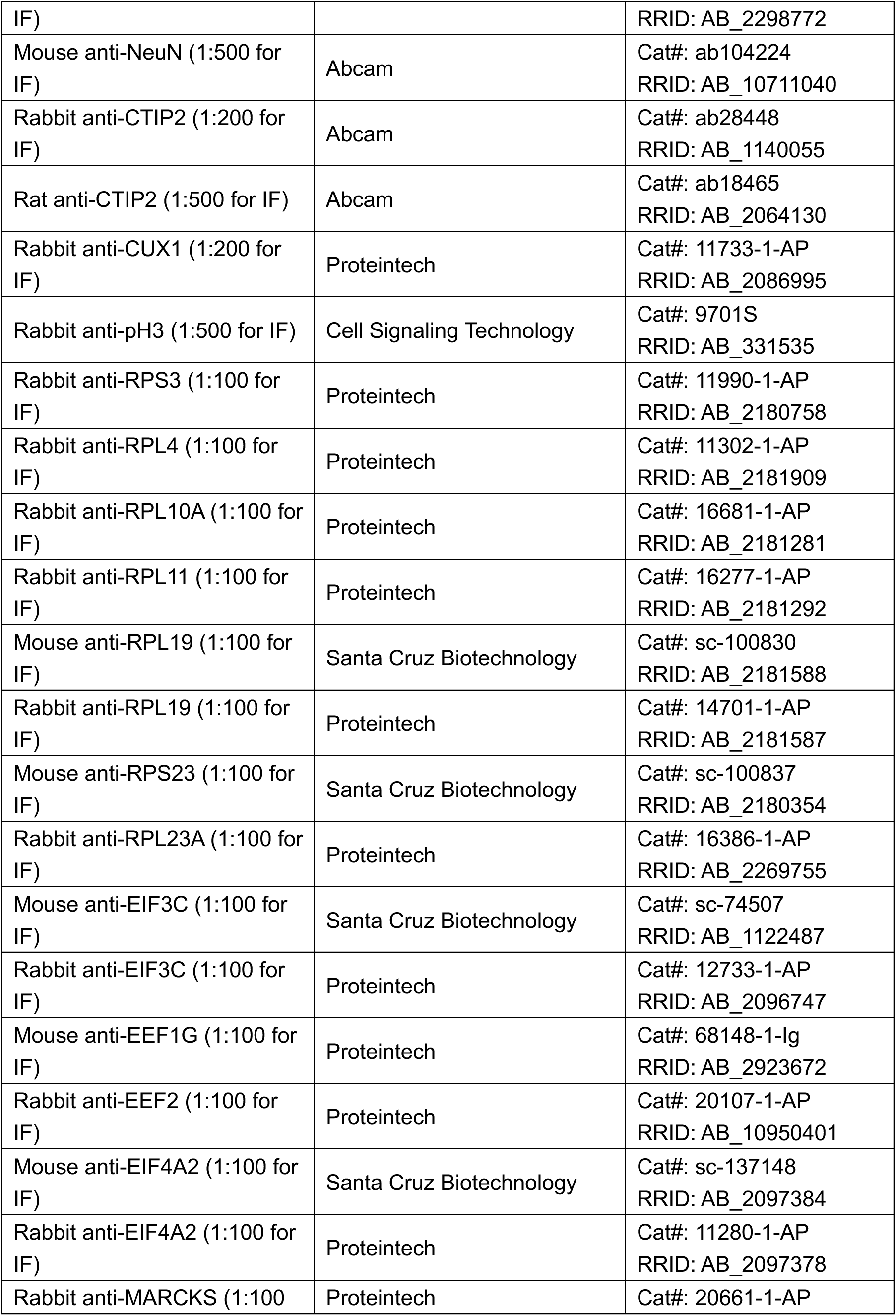
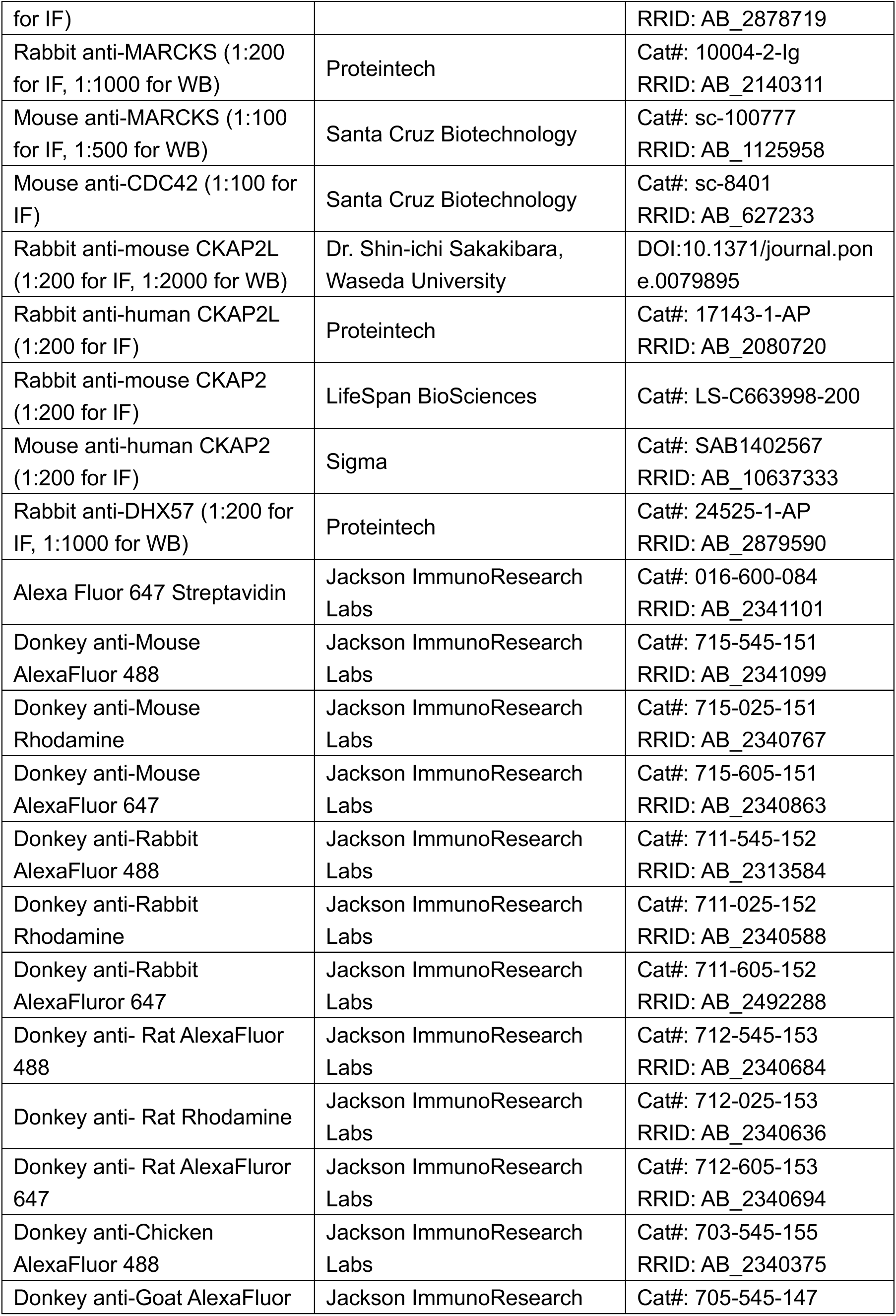
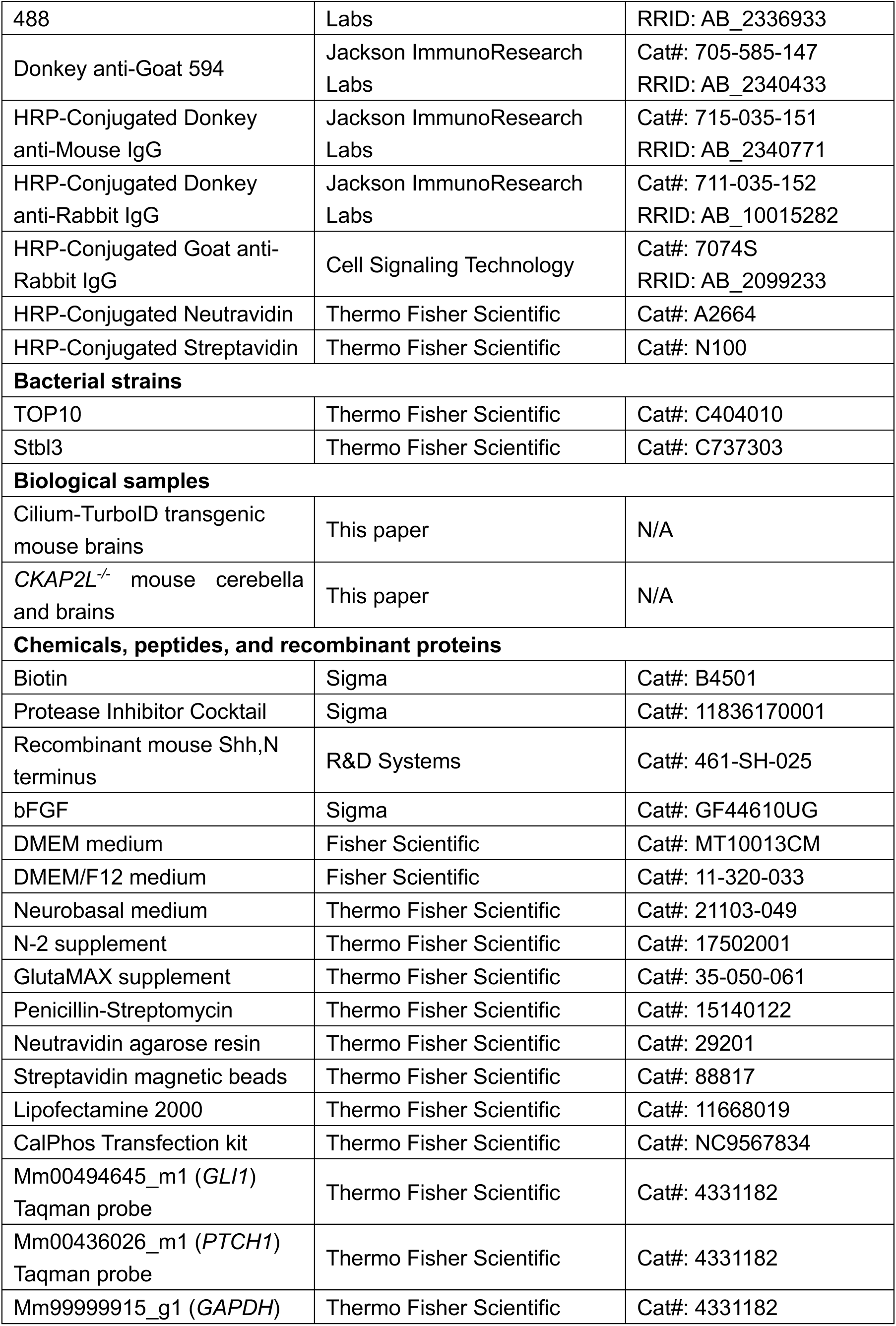
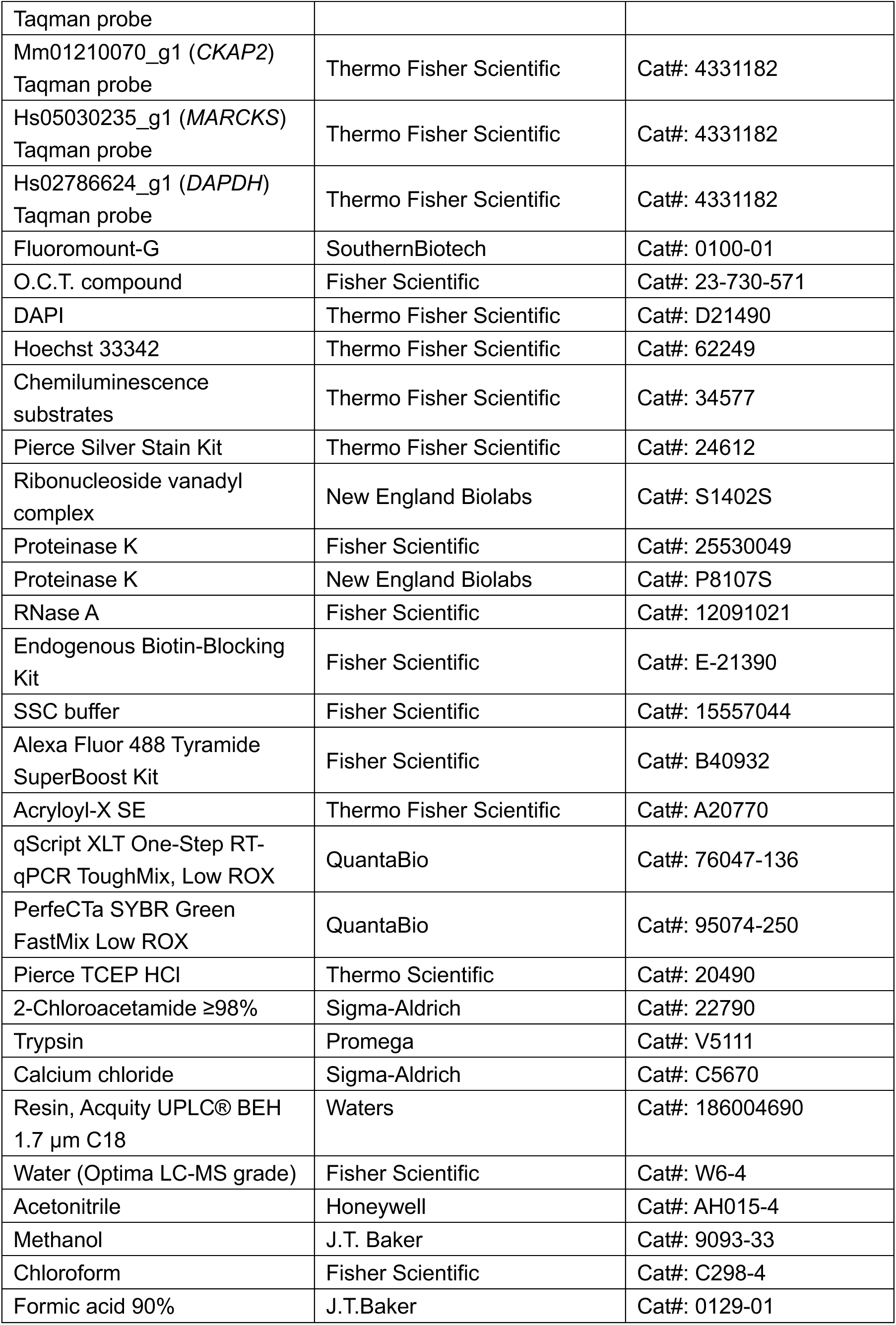
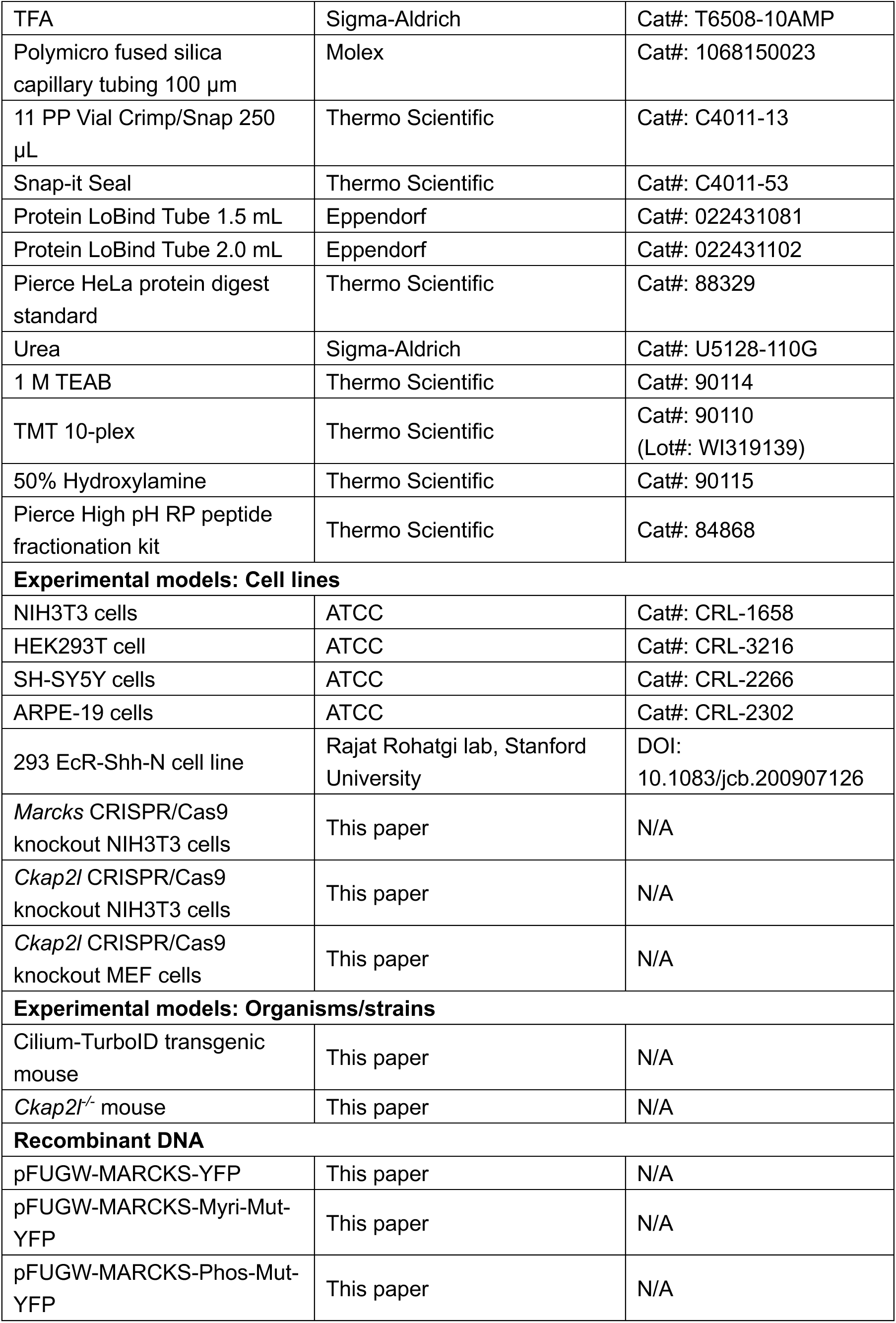
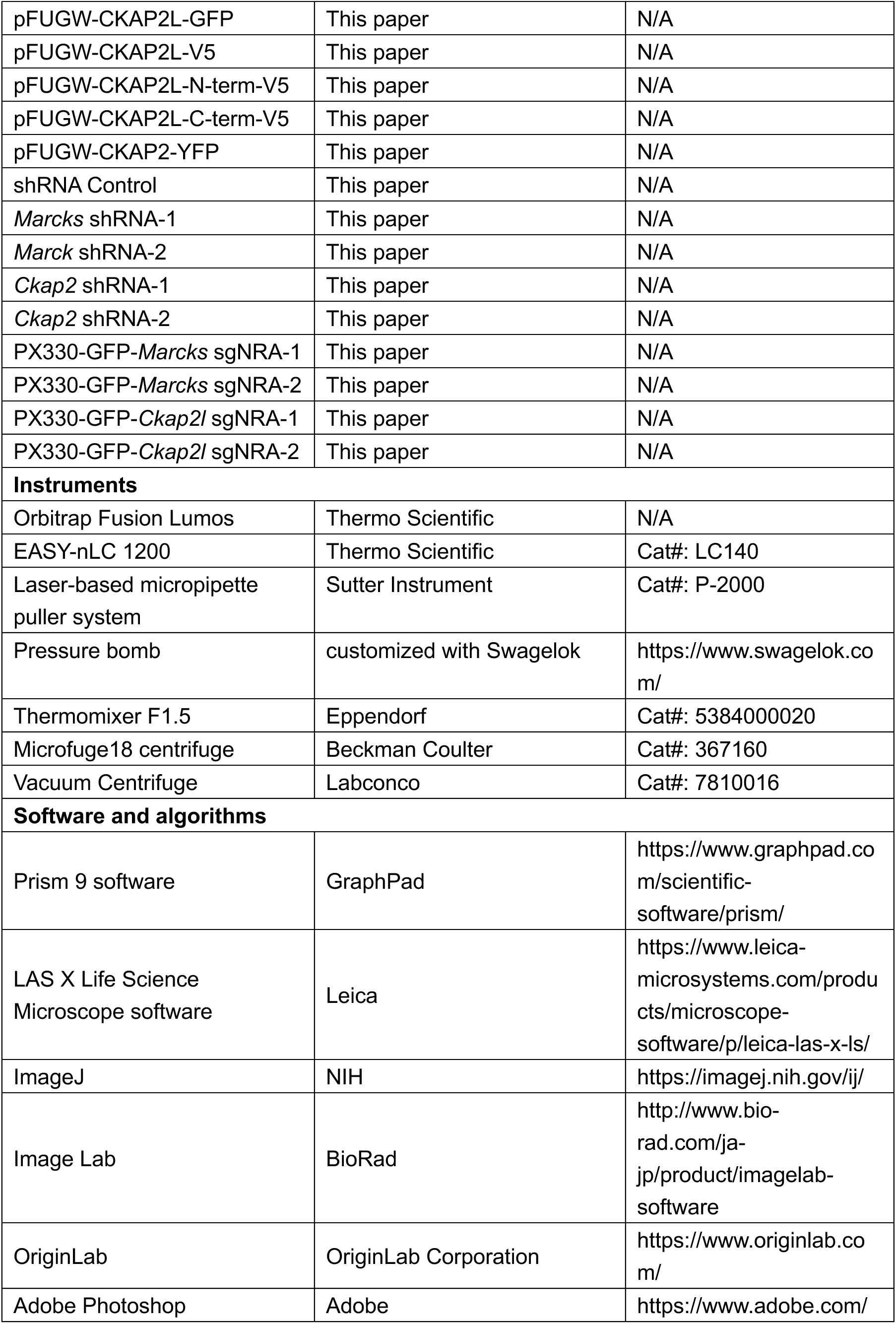
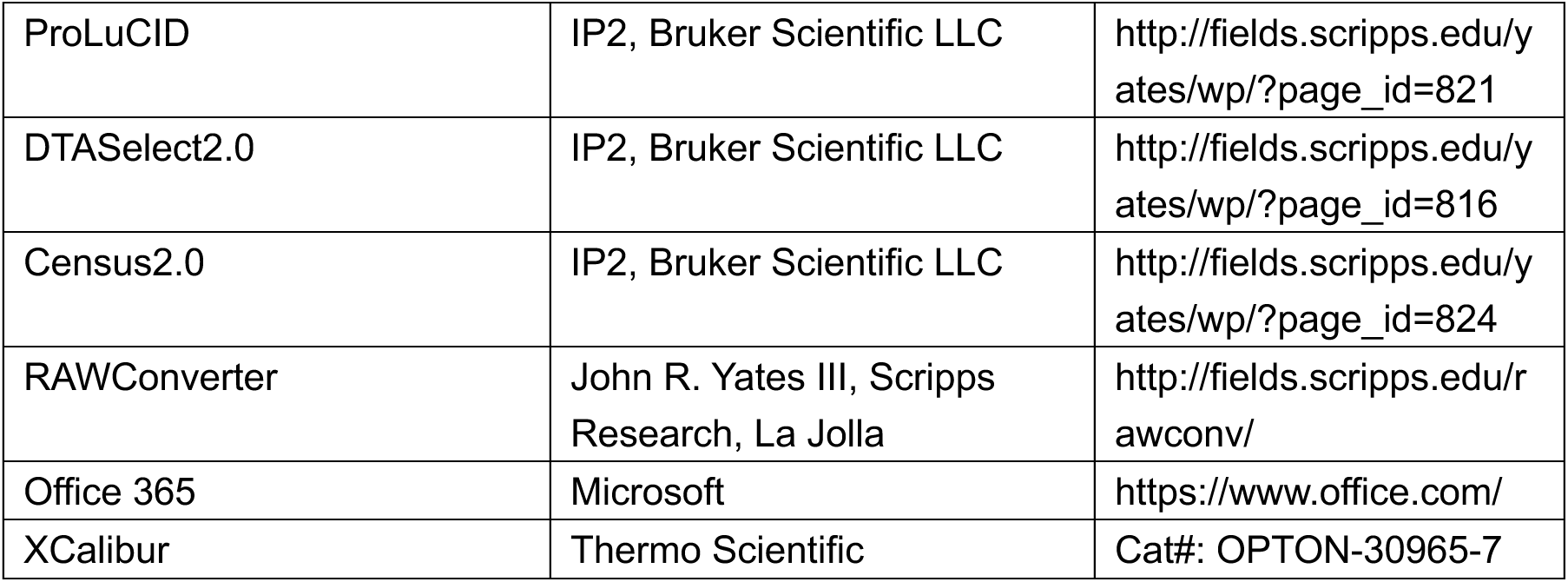

### Mice

All mouse work was performed according to guidelines approved by the IACUC of the University of California, Merced (animal protocol no. 2023-1151). All mice were maintained in a specific pathogen-free animal facility. Mice were monitored daily and housed in a 12 h light / 12 h dark cycle, under the temperature of 66-72°F with 40-60% humidity. The *BLBP*-driven detlaARL13B-GFP-TurboID transgenic mice were generated in C57BL/6 mice by a PiggyBac strategy. *Ckap2l^-/-^* mice were generated in C57BL/6 mice using CRISPR/Cas9, as described in Figure S8A. Embryos were analyzed at the indicated stage described in the figure legends. Adult mice were euthanized by cervical dislocation.

### Primary culture of Radial glia cells

RG were isolated from cortexes of E12.5 embryos. Briefly, isolated cortexes were cut into small pieces and treated with Papain/DNase I. After trituration, single-cell suspensions were harvested by centrifugation and then resuspended in plating medium (Neurobasal medium supplemented with N2, Glutamax, penicillin/streptomycin and 10% Horse serum) for 4-6hr until cells adhered to the bottom of the culture dish. Then, the plating medium was changed into the culture medium (Neurobasal medium supplemented with N2, Glutamax, penicillin/streptomycin, and bFGF).

### Cell lines

NIH3T3 cells (Cat#: CRL-1658), HEK293T cells (Cat#: CRL-3216), SH-SY5Y cells (Cat#: CRL-2266) and ARPE-19 cells (Cat#: CRL-2302) were purchased from ATCC. NIH3T3 cells were cultured in DMEM supplemented with 10% v/v calf serum, and ciliation was induced by reducing the growth media to 0.5% v/v calf serum for 24 h. HEK293T cells were cultured in DMEM supplemented with 10% v/v fetal bovine serum. SH-SY5Y cells were cultured in DMEM/F12 medium supplemented with 10% v/v fetal bovine serum, and ciliation was induced by reducing the growth media to 0.25% v/v fetal bovine serum for 24-36 h. ARPE-19 cells were cultured in DMEM/F12 medium supplemented with 10% v/v fetal bovine serum, and ciliation was induced by DMEM/F12 medium without fetal bovine serum for 48 h. The HEK293T EcR-Shh-N cell line is a gift from Dr. Rajat Rohatgi lab, Stanford University. To induce Shh signaling, growth media were added with ShhN conditioned medium (20-30% v/v, optimal dilution varies batch-wise) produced from HEK293T EcR-Shh-N cells.

### Generation of *Marcks* knockout NIH3T3 cells

*Marcks^-/-^* NIH3T3 cells were generated using CRISPR/Cas9-mediated genome editing with two guide RNAs: #1 (5’-AGGGAGAAGCCACCGCCGAG-3’) and #2 (5’ -ATTTGCTTTGGAAGGCGACG-3’). The guide RNAs were selected according to the online tool Benchling (https://benchling.com/editor), and their sequences were cloned into the PX330-GFP plasmid, respectively. Cell line clones were obtained by single-cell sorting, and *Marcks* deletions were confirmed by immunofluorescence (IF) and Western blot with anti-MARCKS antibodies. Two independent CRISPR/Cas9 knockout cell clones were chosen for the analysis.

### Generation of *CKAP2L* knockout NIH3T3 cells

*Ckap2l^-/-^* NIH3T3 cells were generated using the same method as generating *Marcks* knockout NIH3T3 cells. The two guide RNAs are #1 (5’ -GTCGCCGTAGAGCCATGGTG-3’) and #2 (5’-CCGCGGCTGCGCTGGCAGTG-3’).

### TurboID labeling experiments and subcellular fractionation

TurboID labeling experiments were performed as described previously^36^. Briefly, the freshly dissected dorsal or ventral brains were incubated in the Neurobasal medium in the presence of 50 μM biotin for 15 min. For non-labelled brain tissues, water was added instead of biotin. After 15 min of incubation at 37℃, the medium was removed quickly, and tissues were washed four times with 1× PBS and kept on ice. For immunofluorescence microscopy, tissues were immediately fixed in 4% PFA in PBS. For western blot and proteomic analysis, brain tissues were lysed by subcellular fractionation buffer (20 mM HEPES, 10 mM KCI, 2 mM MgCl_2_, 1 mM EDTA, 1 mM EGTA, 1 mM DTT, pH 7.5 and protease inhibitors) and passed through a 27-G needle 15 times. Nuclei and mitochondria were separated from the supernatant by two sequential centrifugations at 3,000 rpm (720 x g) and 8,000 rpm (10,000 x g). The supernatant was then lysed in lysis buffer (0.5% NP-40, 0.1% SDS, 0.5% sodium deoxycholate, 150 mM NaCl, 50 mM Triethylammonium bicarbonate (TEAB), pH 7.5, and protease inhibitors), sonicated and finally clarified by centrifugation.

### Enrichment of biotinylated proteins

For neutravidin capture, the lysates from the TurboID labeling experiments were measured to determine the protein concentrations and were then adjusted to equal concentrations and volumes as starting material. The lysates were added to washed and equilibrated neutravidin agarose resins, and biotinylated proteins were allowed to bind for 1.5 h at room temperature. Neutravidin agarose resins with bound proteins were washed 2 times with lysis buffer and 4 times with 1 x PBS. For Western blot analysis, the beads were eluted with 2 x SDS sample buffer.

Streptavidin capture was performed as described previously^36^. Briefly, the lysates were added onto washed and equilibrated streptavidin magnetic beads, and biotinylated proteins were then allowed to bind for 1.5 h at room temperature. Streptavidin beads with bound proteins were washed extensively with a series of buffers: 2 times with lysis buffer, 1 time with 1 M KCl, 1 time with 0.1 M Na_2_CO_3_, 1 time with 2 M Urea in 10 mM Tris-HCl, pH 8.0, 2 times with lysis buffer, and finally 2 times with 1 x PBS. For Western blot analysis, the beads were eluted with 2 x SDS sample buffer.

### On-beads trypsin digestion and TMT labeling

Enriched proteins bound to beads were denatured with 8 M urea in 100 mM TEAB, disulfide bonds reduced with 5 mM TCEP and free cysteines alkylated with 2-Chloroacetamide. Denatured proteins bound to beads were precipitated with methanol-chloroform and digested with trypsin in 1 M urea and 100 mM TEAB. Resulting peptides were dried by vacuum centrifugation and then labeled with 10-plex TMT as described in the manufacturer’s protocol. A reference TMT channel was used for internal normalization using a tenth of all protein in other 9 channels. TMT-labeled peptides were pooled, dried and reverse-phase fractionated into 4 parts at high pH using 12.5%, 17.5%, 22.5% and 50% acetonitrile, as per manufacturer’s protocol.

### Mass spectrometry

The samples were analyzed on a Fusion Lumos mass spectrometer. Samples were injected directly onto a 25 cm, 100 μm ID column packed with BEH 1.7 μm C18 resin. Samples were separated at a flow rate of 300 nL/min on a nLC 1200. Buffer A and B were 0.1% formic acid in 5% and 80% acetonitrile, respectively. A gradient of 1–25% B over 100 min, an increase to 40% B over 20 min, an increase to 90% B over another 10 min and held at 90% B for a final 10 min of washing was used for 140 min total run time. Column was re-equilibrated with 15 μL of buffer A prior to the injection of sample. Peptides were eluted directly from the tip of the column and nanosprayed directly into the mass spectrometer by application of 2.8 kV voltage at the back of the column. The Lumos was operated in a data dependent mode. Full MS1 scans were collected in the Orbitrap at 120k resolution. The cycle time was set to 3 s, and within this 3 s the most abundant ions per scan were selected for CID MS/MS in the ion trap. MS3 analysis with multinotch isolation (SPS3) was utilized for detection of TMT reporter ions at 7.5k resolution. Monoisotopic precursor selection was enabled and dynamic exclusion was used with exclusion duration of 10 s.

### MS data processing

Protein and peptide identification was done with Integrated Proteomics Pipeline (IP2, Bruker Scientific LLC). Tandem mass spectra were extracted from raw files using RawConverter^99^, and searched with ProLuCID^100^ against a database comprising UniProt reviewed (Swiss-Prot) proteome for *Mus musculus* (UP000000589) with modified Arl13b (UniProt, Q640N2; amino acids 1 and 19-356 deleted, EGFP-TurboID fused at C-terminus), Streptavidin (UniProt, P22629) added, and a list of general protein contaminants. The search space included semitryptic peptide specificity with unlimited missed cleavages. Carbamidomethylation (+57.02146 C) and TMT (+229.1629 K and N-terminus) were considered static modifications. Data was searched with 50 ppm precursor ion tolerance and 500 ppm fragment ion tolerance. Identified proteins were filtered using DTASelect2^101^ and utilizing a target-decoy database search strategy^102^ to limit the false discovery rate to 1%, at the spectrum level. A minimum of one peptide per protein and one tryptic end per peptide were required, and precursor delta mass cutoff was fixed at 5 ppm. Statistical model for tryptic peptides (trypstat) was applied. Census2 isobaric-labeling analysis was performed based on the TMT reporter ion intensity using default parameters, thresholding isobaric purity at 0.6 and normalizing to the internal reference channel^103^.

### GO analysis

Gene ontology (GO) enrichment analysis of biological processes was plotted according to the rich factor in Metascape (https://metascape.org/gp/#/main/step1). The top 20 enriched biological processes are represented in the scatter plot.

### Whole-mount dissection and immunofluorescence

The E12.5 brains were harvested and fixed in 4% (w/v) paraformaldehyde (PFA) in 1x PBS for 2-12 h at 4°C. The brains were then rinsed 2 times with PBS and further dissected into dorsal and ventral regions. Prior to staining, the brain tissues were permeabilized with 0.2% Triton X-100 in PBS for 10min and blocked for 1 h in blocking buffer: 2% donkey serum in PBS. The brain tissues were incubated with primary antibody overnight at 4℃, washed 3 times with PBS and incubated with secondary antibody for 1h, followed by DAPI staining for 10 min at room temperature. The brain tissues were then dissected into small pieces before mounting with Fluoromount-G.

### Cryostat sectioning and immunofluorescence

The dissected brains were fixed in 4% PFA in PBS overnight at 4°C and then incubated in 30% sucrose overnight or until they sunk to the bottom. The brains were embedded in O.C.T. compound and sliced into 10-20um thick sections with Leica CM1850-3-1 Cryostat Microtome. For immunofluorescence on cryosections, brain slices were blocked for 1 h at room temperature in the blocking solution (2% goat serum, 0.2% Triton X-100 in PBS). Subsequently, the sections were incubated with primary antibodies for 2 h at room temperature or 12 h at 4°C, washed 3 times with PBS and incubated with secondary antibodies for 1h, followed by DAPI staining for 10 min at room temperature. Last, sections were mounted in Fluoromount-G for imaging.

### SDS-PAGE and Western blot

Cells were harvested and lysed in RIPA buffer (1% NP-40, 0.1% SDS, 0.5% sodium deoxycholate, 150 mM NaCl, 25 mM Tris/HCl, pH 7.5, and protease inhibitors). Lysates were clarified by centrifugation (16,000 ×g at 4°C for 15 min), and 20-30 µg protein was separated on SDS-PAGE gels and transferred onto PVDF membranes. Membranes were blocked with 5% BSA, incubated and washed with primary and secondary antibodies. The protein bands were detected with chemiluminescence substrates, and images were captured by the Biorad ChemiDoc system.

### Silver staining

The SDS-PAGE gel was washed in ultrapure water, fixed in 30% ethanol:10% acetic acid solution, and washed with 10% ethanol solution and ultrapure water. The gel was then incubated in a sensitizer working solution for exactly 1 minute and washed with ultrapure water, followed by incubating in a stain working solution for 30 min. The gel was quickly washed with ultrapure water, immediately added developer working solution, and incubated until protein bands appeared. Last, the gel was stopped by incubating the stop solution (5% acetic acid).

### RNA-FISH-combined immunofluorescence (IF) staining

When RNA FISH is combined with IF, we performed IF (under RNase-free conditions) before FISH. Briefly, the freshly PFA-fixed samples were permeabilized for 5min in 0.2 % v/v Triton X-100 in PBS containing 2 mM ribonucleoside vanadyl complex (RVC), followed by proteinase K treatment and mock RNA digestion. The samples were then blocked with the blocking buffer, incubated with primary and secondary antibodies, and fixed again. Before RNA FISH, the endogenous biotin or biotinylated proteins in the samples were blocked using the Endogenous Biotin-Blocking Kit. The samples were pre-hybridized for 1-2 hours at 42°C, then hybridized overnight with the biotin-conjugated Oligo(dT)20 probe in hybridization buffer and washed with SSC buffer. The samples then went through signal amplification with the Alexa Fluor 488 Tyramide Super Boost Kit. Briefly, samples were incubated with HRP-conjugated streptavidin for 30-60 min at room temperature and treated with tyramide working solution for 3 min. Within this time, HRP converted Alexa Fluor 488-conjugated-tyramide into a highly reactive form that covalently links to tyrosine residues on proteins near HRP. The reaction was stopped by the reaction stop reagent. The tissues were then mounted for imaging.

### Lentivirus preparation

To generate lentivirus, HEK293T cells were seeded into a 10 cm dish and later these cells should be > 90% confluent at the time of transfection. The cells were transfected with 15 μg lentiviral plasmid, 10 μg psPAX2 plasmid and 5 μg pMD2.G plasmid. 48 h after transfection, the supernatant containing lentivirus was harvested and concentrated with the concentrator solution. The aliquoted virus was then frozen at -80°C.

### Immunofluorescence staining on cells

Cells were fixed in 4% PFA for 10 min, permeabilized and blocked with blocking buffer (2% donkey serum, 0.2% triton X-100, in PBS) for 1 h at room temperature. Cells were then incubated with primary antibodies for 1 h at room temperature or 4°C overnight and incubated with secondary antibodies for 1 h, followed by staining in DAPI for 10 min. Finally, cells were mounted on glass slides using Fluoromount G.

### Expansion microscopy

Fixed cells with 4% PFA were incubated in primary antibodies and then were fixed in 4% PFA for 20 min. After fixation, cells were incubated in the AcX (Acryloyl-X SE) solution for 2 to 3 h and then polymerized in the gelling solution. The gel was digested with proteinase K for 12 h at room temperature. After digestion, the gel was incubated in the secondary antibody for 2 h at room temperature. The gel was then washed, expanded by four washes with excessive water and mounted on glass slides with superglue.

### Quantitative real-time PCR

Total RNAs were extracted using the Trizol reagent. Quantitative PCR was performed in Quantstudio 3 System (Applied Biosystems) with 100 ng total RNA per reaction and qPCR reagents (qScript XLT One-Step RT-qPCR ToughMix, Low ROX). The TaqMan gene expression probes used were Hs05030235_g1 (*MARCKS*), Hs02786624_g1 (*GAPDH*), Mm01210070_g1 (*CKAP2*), Mm00494645_m1 (*GLI1*), Mm00436026_m1 (*PTCH1*) and Mm99999915_g1 (*GAPDH*) to normalize the samples.

For the genotyping analysis of the *detlaARL13B-GFP-TurboID* transgenic mice, quantitative PCR was performed in Quantstudio 3 System with the qPCR reagents (PerfeCTa SYBR Green FastMix Low ROX). The following primers were used for qRT-PCR: transgene (F: 5’-GAATCGGCGAGCTGAAGAGT-3’ and R: 5’-CATTGGGCCATTTGACTCGC-3’), *GAPDH* (F: 5’-AGCCATCAGCTATGCACGTA-3’ and R: 5’-GCCTCGGCTGCTCAAAGAAT-3’). Transcript levels were calculated relative to *GAPDH* using the ΔΔCt method.

### Image acquisition

Confocal images were acquired on an LSM880 confocal microscope (Zeiss) and an Olympus IX83 microscope (Evident) equipped with a spinning disk confocal unit (CSU–W1; Andor). Immunofluorescence images of cell lines were imaged on a Leica DMi8 (LAS X software) with Plan Apochromat oil objectives (63x, 1.4 NA). For expansion microscopy, imaging was performed by an AiryScan LSM880 confocal microscope (Zeiss) with a 63x lens. Widefield fluorescence images were acquired on a Leica Mica confocal microscope with a 10x lens. Western blot and silver staining images were acquired on a ChemiDoc MP Imaging System (Bio-Rad).

## QUANTIFICATION AND STATISTICAL ANALYSIS

### Quantification

For all analyses, the number of replicates is listed in the figure legends. ImageJ software was used for the measurement of brain width, cortex length, axis length, area of cerebella, and cell number. For quantification on cultured cells, 5-10 images for each condition were randomly taken from the coverslips. For cilium quantification, 50-100 cilia from the 5-10 images were counted. Cilium length and protein intensity were measured in Leica LAS X software. For mutant mice, 10 adult mice were used for the measurement of brain size. For quantification of embryonic and postnatal cortex thickness, 6-7 brains were used for each condition. For western blot, quantitation of bands was performed in ImageJ.

### Statistical analysis

All statistical tests were performed using GraphPad Prism, and data are presented as means ± SD. All statistical methods are listed in the figure legends. Statistical tests were performed using two–way analysis of variance (ANOVA) with Tukey’s multiple comparisons for quantification of ciliary Smo and Gli2 intensity in the cilia and quantitative real-time PCR, and using one-way ANOVA followed by the multiple comparisons (Tukey test) for the analysis of adult mouse brains and quantification of the percentage of ciliated cells, cilia length, *MARCKS* and *CKAP2* RNA levels. For the analysis of normal distribution in other tests and volcano plot graphs, we used a two-tailed unpaired Student’s t-test. For all analyses, statistically significant data are indicated as: *p < 0.05, **p < 0.01, ***p < 0.001, ****p < 0.0001. Non-significant data is indicated as ns.

## SUPPLEMENTAL INFORMATION

Supplemental information can be found in this paper.

## ACKNOWLEDGMENTS

We thank Dr. Rajat Rohatgi (Stanford University) for the gifts of CRISPR/Cas9 reagents, and Dr. Matthew Scott lab for the antibody against SMO. We thank Dr. Shin-ichi Sakakibara (Waseda University) for the antibody against mouse CKAP2L. We thank Dr. David Gravano (Stem Cell Instrumentation Foundry, University of California, Merced) for fluorescence-activated cell sorting. TurboID is provided by Dr. Alice Ting at Stanford University. The data in this work was collected, in part, with a confocal microscope acquired through the National Science Foundation MRI Award Number DMR-1625733. O.T. Gutierrez and Y. Al-Issa are graduate student fellow of NSF-CREST Center for Cellular and Biomolecular Machines at the University of California, Merced (NSF-HRD-1547848 and NSF-EES-2112675). Research in the laboratory of X.G. was supported by NIH/NCI (CA235749), NIH/NIGMS(GM143276), NIH/NCI (CA274595), and NSF CAREER award (IOS-2143711). Research from J.R.Y. laboratory was supported by NIH/NIGMS P41GM103533.

## AUTHOR CONTRIBUTIONS

X. Ge and X. Liu conceived the project and designed the experiments. X. Liu performed, analyzed and interpreted most of the data on mouse and cell experiments. O.T. Gutierrez participated in harvesting the embryonic brain tissues for mass spectrometry. G. Kaur participated in the maintenance of mouse strains and assisted in immunohistology experiments. Y. Al-Issa participated in characterizing the ciliary localization of CKAP2L and its truncates. S. Baboo and J.K. Diedrich performed on-beads digestion and TMT mass spectrometry under the supervision of J.R. Yates. O.T. Gutierrez and E. Cai characterized the CRISPR/Cas9 knockout cell lines. X. Ge and X. Liu wrote the paper.

---

**Figure S1.**
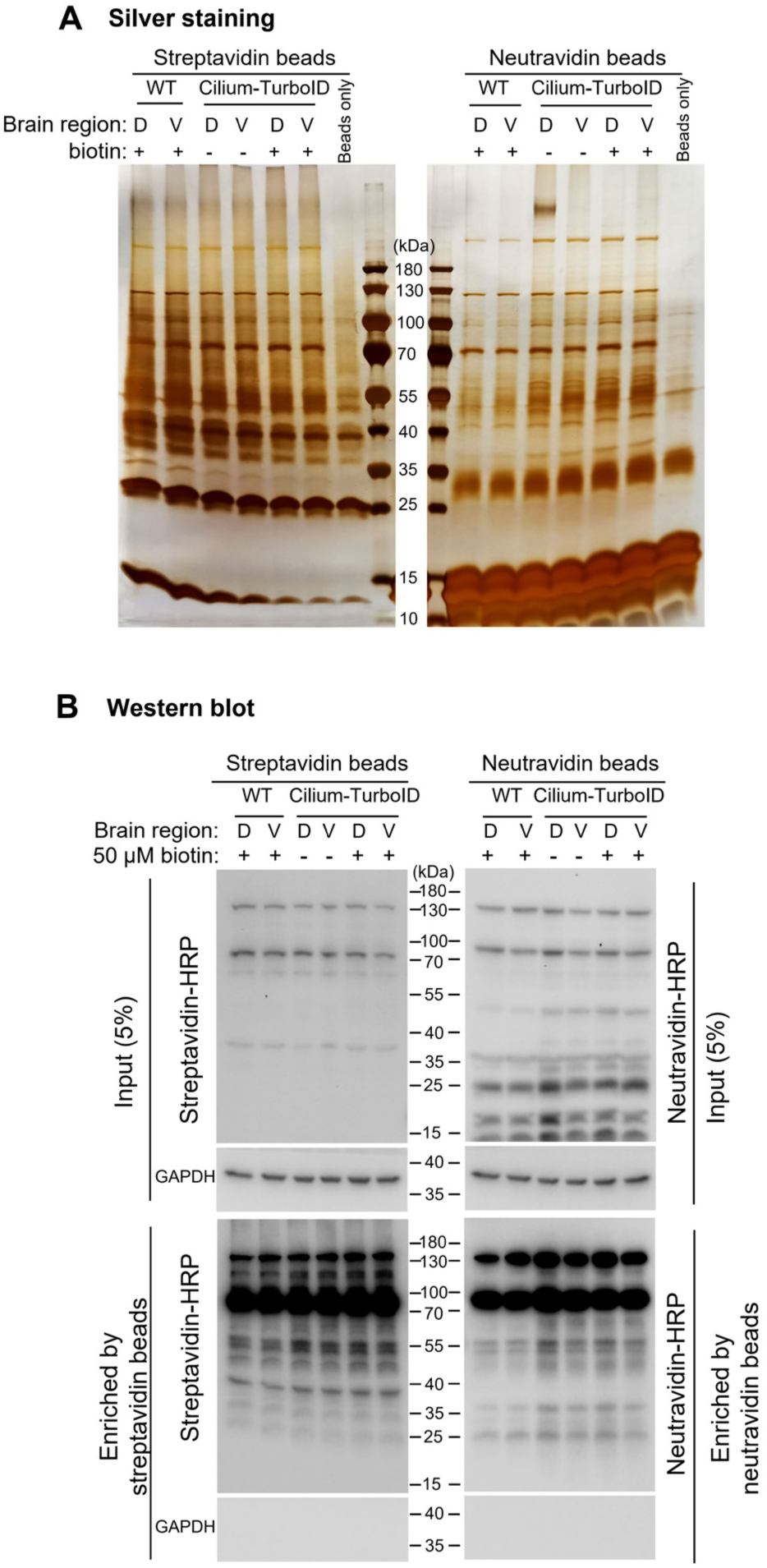
Purification of biotinylated proteins from the brain of Cilium-TurboID transgene mice. (**A**) Silver staining analysis of proteins enriched by streptavidin and neutravidin beads. The dorsal (D) and ventral (V) brain regions from E12.5 WT or Cilium-TurboID mice were incubated with or without 50 µM biotin for 15min. The tissues were then lysed and biotinylated proteins were captured by streptavidin or neutravidin beads. In each lane, lysates from 10-12 brains were used. (**B**) Western blotting analysis of biotinylated proteins enriched by streptavidin and neutravidin beads. The biotinylated proteins in brain tissues were processed similarly as in (A). The overall biotinylated proteins were detected by streptavidin-HRP or neutravidin-HRP.

**Figure S2.**
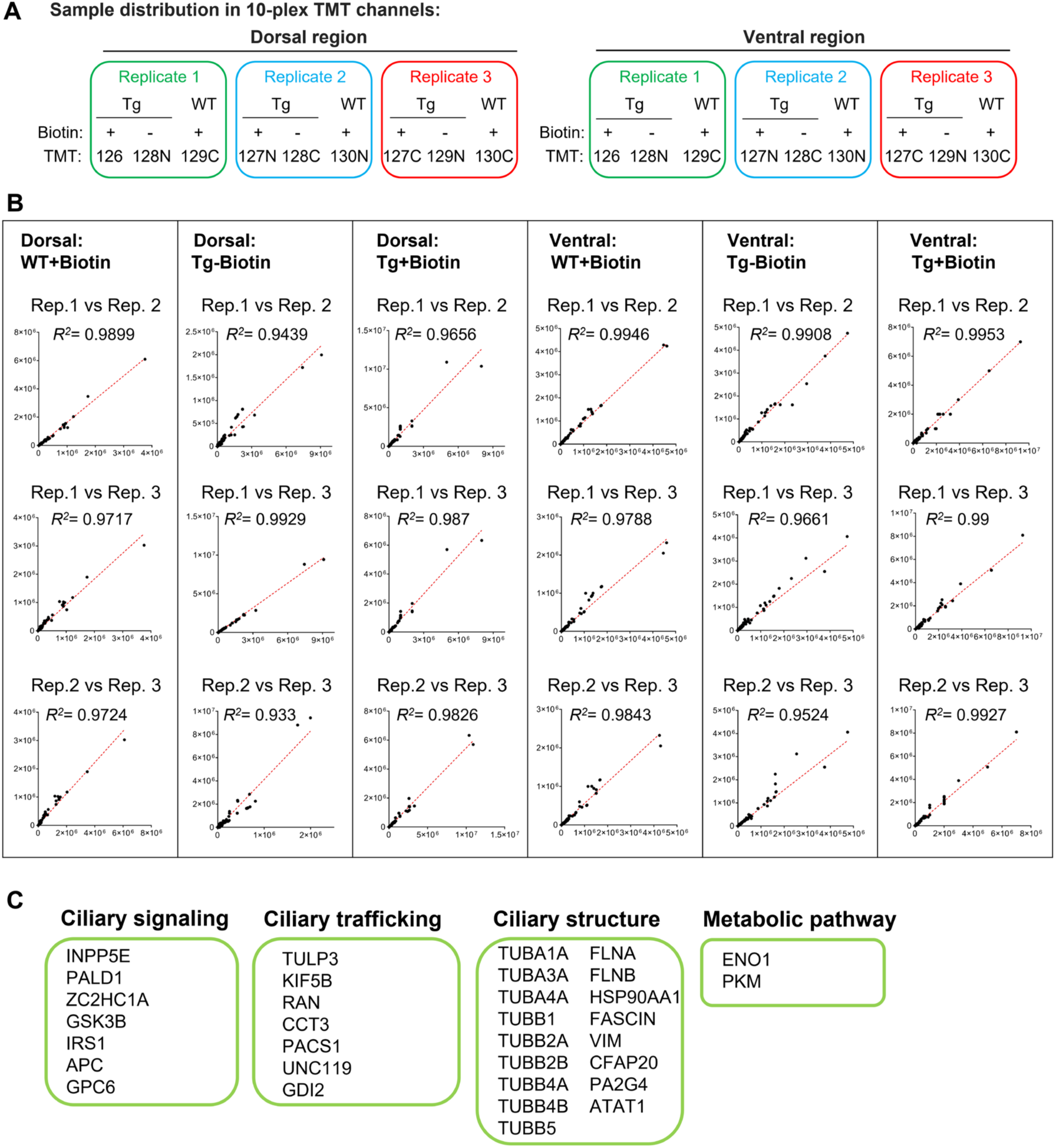
Sample distribution in 10-plex TMT channels for quantitative proteomics and correlation of biological replicates in mass spectrometry. (**A**) Sample distribution in 10-plex TMT channels. Three biological replicates are included for each condition. The dorsal and ventral regions are processed in parallel. Labels in the TMT row indicate the TMT channel used for each condition. (**B**) Correlation of biological replicates for each experimental condition. (**C**) The previously reported ciliary candidates recovered in this study are listed and grouped into functional categories.

**Figure S3.**
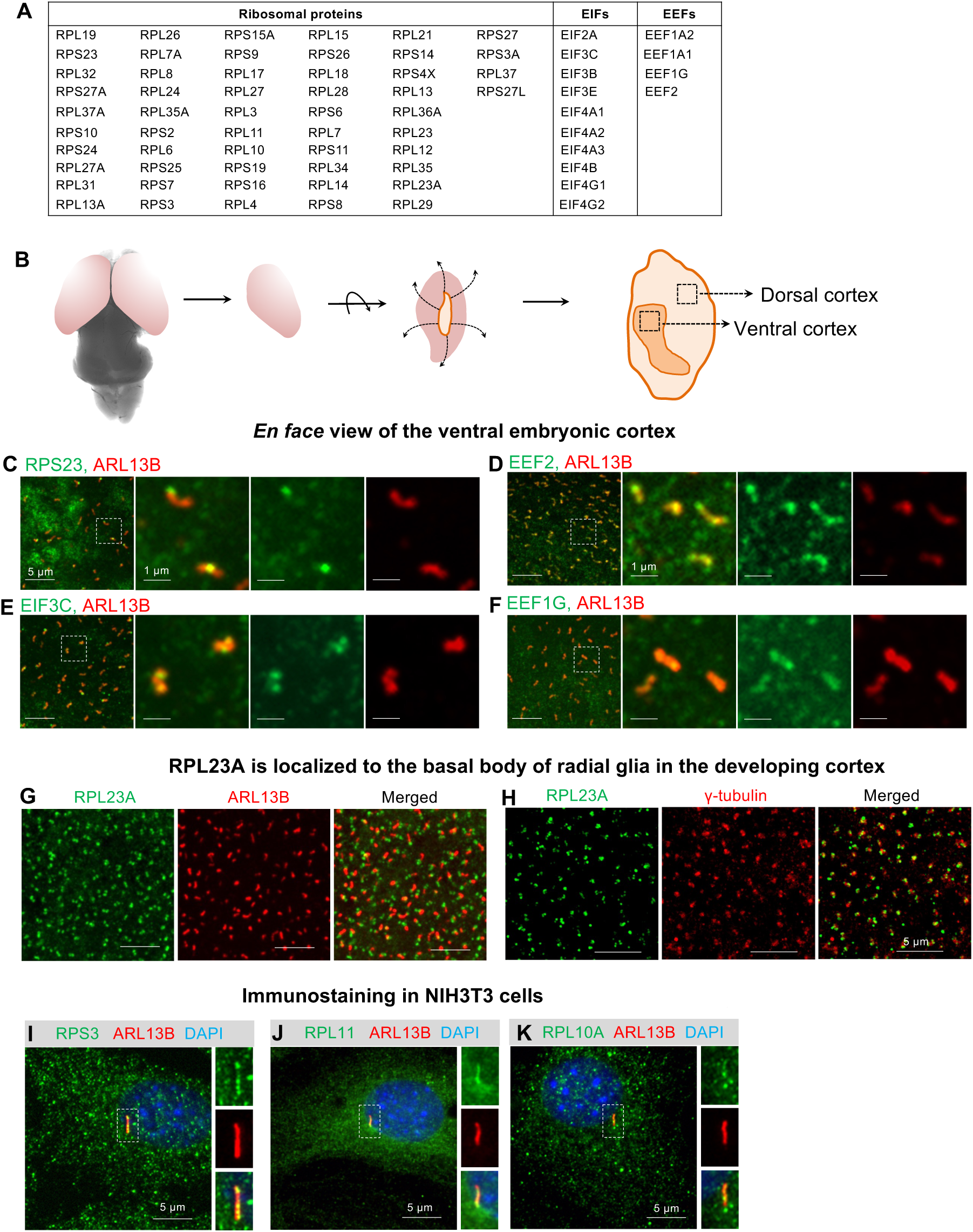
*In vivo* ciliary proteomics uncover translation machinery components in the cilia of radial glia. (**A**) List of ribosome components and translation machinery proteins identified in this study. (**B**) Diagram showing the whole-mount preparation of cortex for *en face* imaging in the dorsal and ventral brain region. (**C**-**F**) *En face* view of the ventral cortex in the whole-mount E12.5 mouse brain. The brain tissues were immuno-stained with antibodies against the indicated translation machinery proteins. Primary cilia are labeled with ARL13B (red). Regions within the white dashed boxes are magnified and shown on the right side. (**G**-**H**) *En face* view of the cortex in the whole-mount E12.5 mouse brain. The brain tissues were immuno-stained with RPL23A, and co-stained with ARL13B or γ-tubulin. (**I**-**K**) Representative images of NIH3T3 cells immunostained with the indicated antibodies. Areas in the white dashed boxes are magnified and displayed at the side.

**Figure S4.**
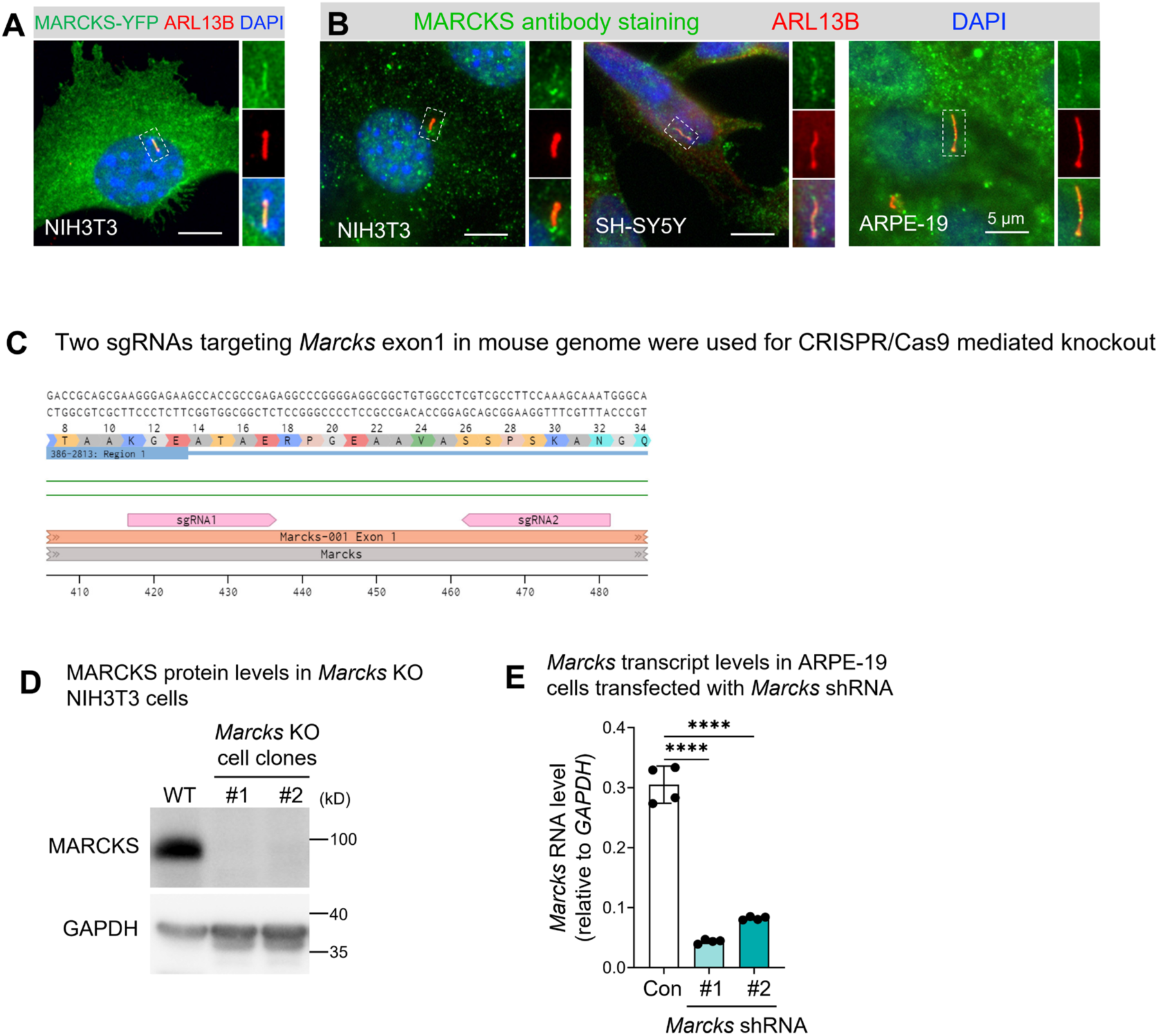
MARCKS localizes in the primary cilia. (**A**) MARCKS-YFP localizes in the primary cilia of NIH3T3 cells. The cilium in the white dashed box is magnified and displayed at the side. (**B**) Immunostaining with anti-MARCKS antibody shows that endogenous MARCKS is present in the cilium of NIH3T3, SH-SY5Y, and ARPE-19 cells. The cilia in the white dashed boxes are magnified and displayed at the side. (**C**) Two guide RNAs were used in CRISPR/Cas9 to target exon 1 of mouse *Marcks*. (**D**) Western blot analysis showing depletion of MARCKS protein in the *Marcks* CRISPR/Cas9 knockout NIH3T3 cell clones. (**E**) *Marcks* shRNA significantly reduced *Marcks* gene expression in ARPE-19 cells. *Marcks* mRNA levels were measured by qPCR. Control (Con) shRNA is a non-targeting scrambled shRNA.

**Figure S5.**
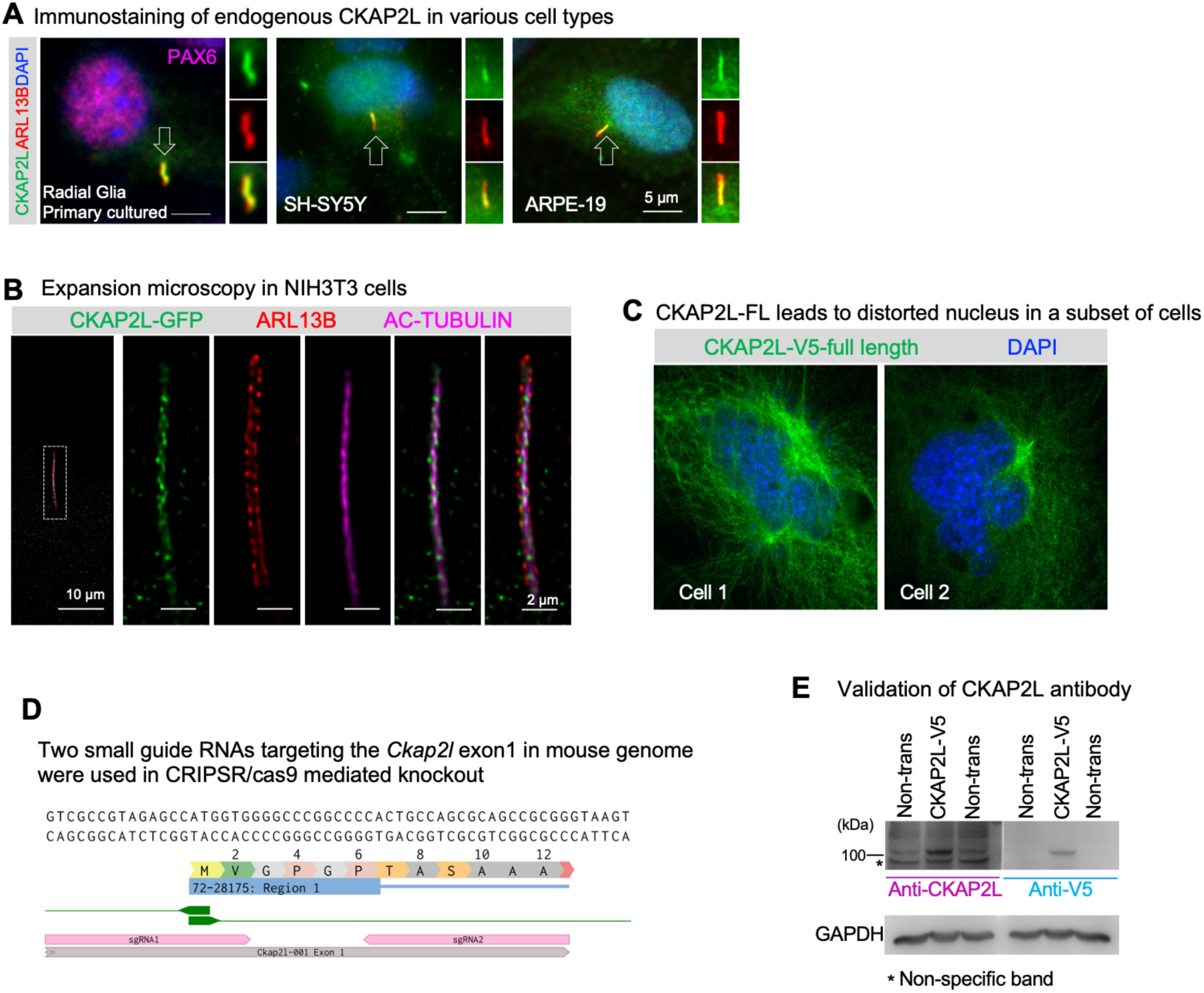
CKAP2L localizes to the primary cilia. (**A**) Immunostaining with anti-CKAP2L antibody shows that endogenous CKAP2L is present in the cilium of primary cultured radial glia, SH-SY5Y, and ARPE-19 cells. The cilia in the white dashed boxes are magnified and displayed at the side. (**B**) NIH3T3 cells was transfected with CKAP2L-GFP plasmid, and subjected to expansion microscopy. ARL13B staining labeled the cilia membrane (red). Ac-Tubulin staining labeled the axoneme (magenta). The CKAP2L signal was detected in the space between the ciliary membrane and the axoneme. (**C**) Representative images showing that over-expressing CKAP2L full length in *Ckap2l* KO cells induced disorganized spindle formation and disorganized nuclei. (**D**) Two guide RNAs were used in CRISPR/Cas9 to target exon 1 of mouse *Ckap2l*. (**E**) Validating CKAP2L antibody used in this study by Western blot. Cell lysate from non-transfected cells or CKAP2L-V5 expressing cells were loaded to the SDS-PAGE and analyzed side by side. Blotting with V5 antibody highlighted one specific band corresponding to the true size of CKAP2L. Asterisk indicates the non-specific band.

**Figure S6.**
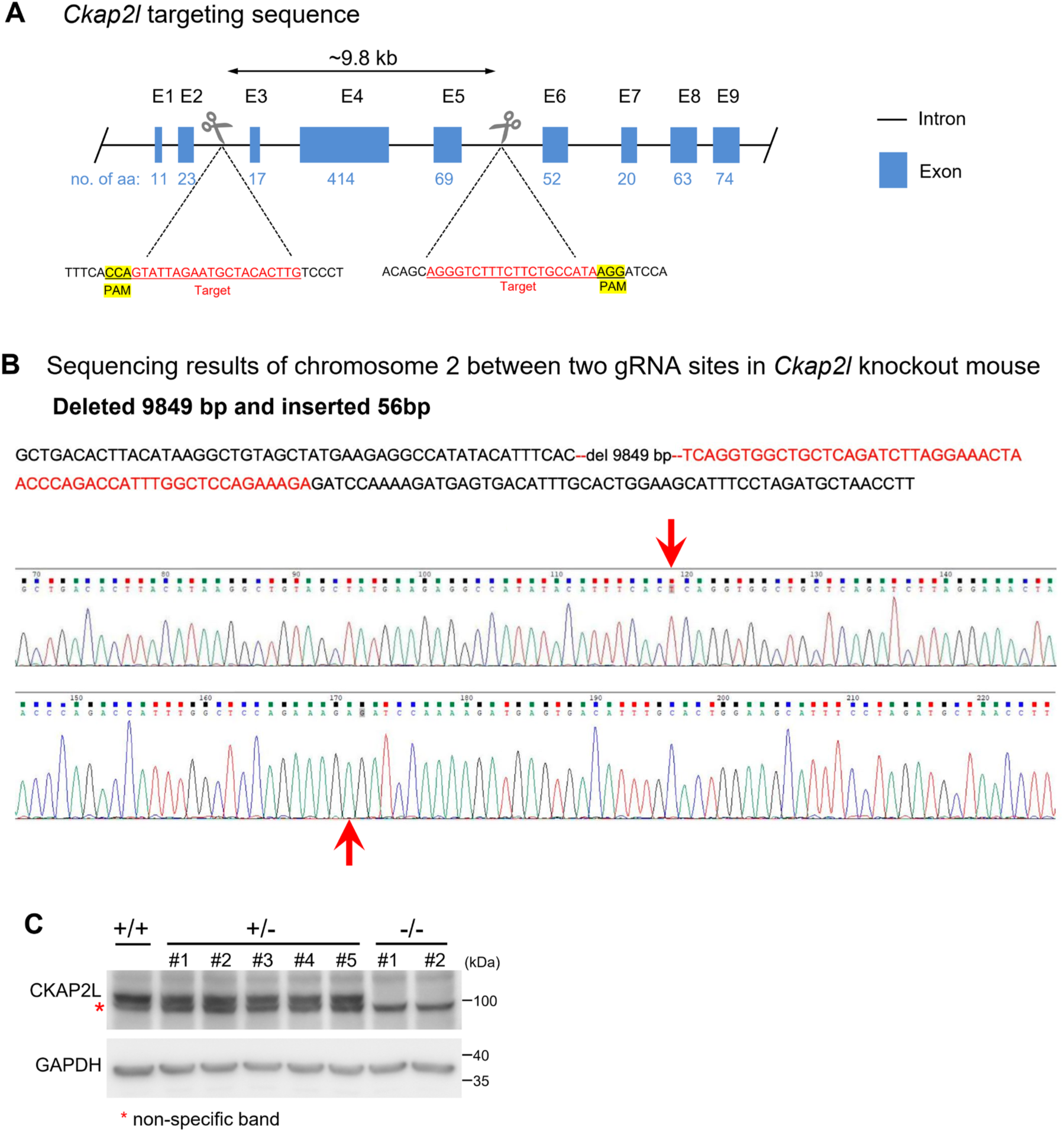
Generation of *Ckap2l* knockout mouse. (**A**) A schematic of the CRISPR/Cas9-mediated *Ckap2l* knockout in chromosome 2 of the mouse genome. Two guide RNAs are used. Sequences of the two sgRNAs are shown in red. The resulting deletion of ∼9.8kb removed exon 3-5 of mouse *Ckap2l*, leading to abolishment of the entire protein. (**B**) Sequencing results of mouse chromosome 2 between two gRNA sites. The 9849bp deletion and 56bp insertion take places between the two red arrows. (**C**) Western blot analysis confirmed the absence of CKAP2L protein in the *Ckap2l^-/-^* mouse brain. Asterisk indicates a non-specific band.

**Figure S7.**
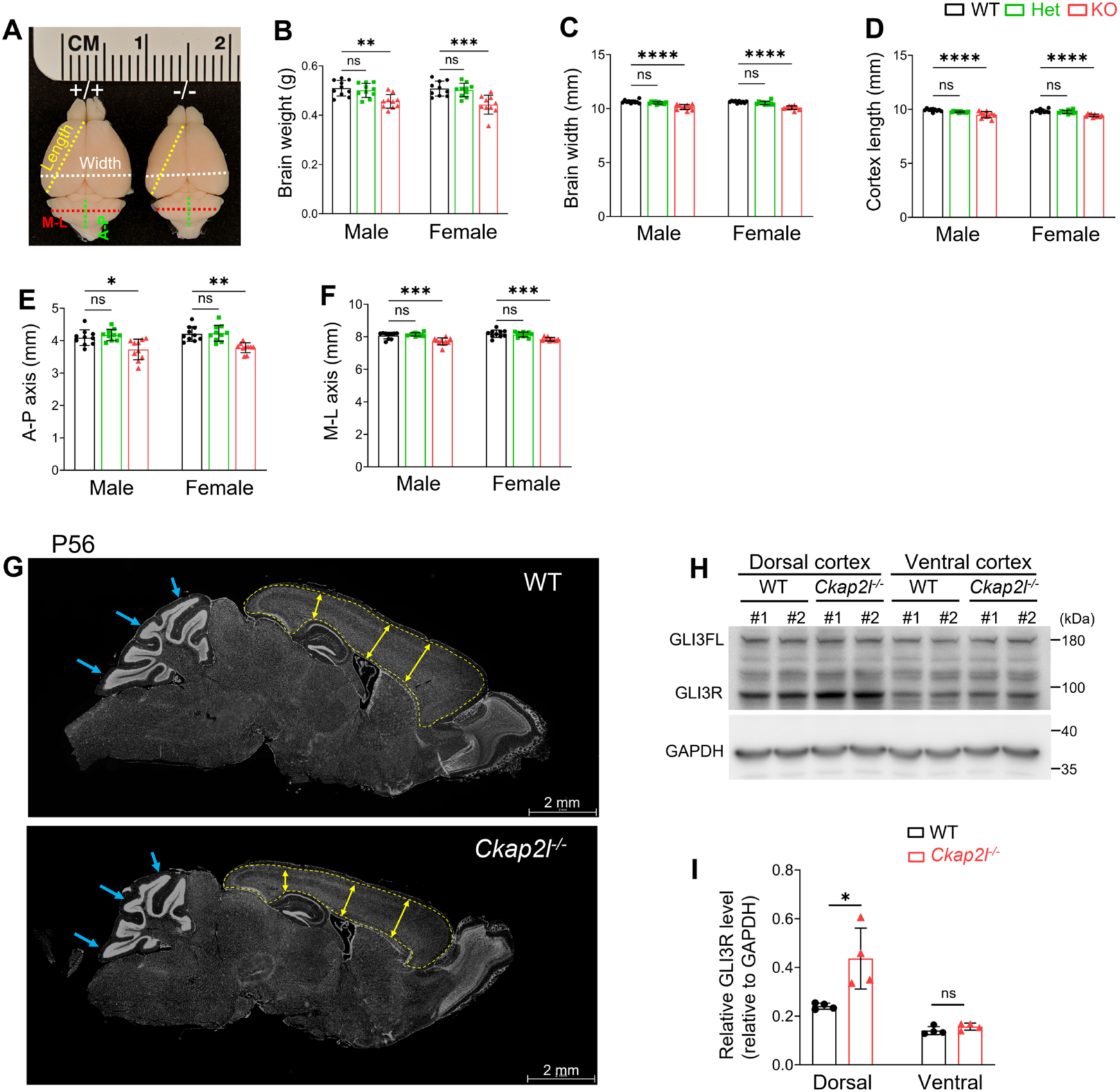
*Ckap2l* loss reduces the size of the brain. (**A**) Brain images of adult (P56) WT and *Ckap2l^-/-^* mice. A-P, anterior-posterior; M-L, medial–lateral. (**B**-**F**) Adult P56 *Ckap2l^-/-^* mice exhibited reduced brain weight (**B**), brain width (**C**), cortex length (**D**), cerebellar anterior-posterior (A-P) (**E**) and medial–lateral (M-L) (**F**) axis lengths. n = 10 mice per group. Data are presented as mean ± SD. Statistics: One-way ANOVA with multiple comparisons (Tukey test). ****, p < 0.0001; ***, p < 0.001; **, p < 0.01; *, p < 0.05; ns, not significant. (**G**) Coronal sections of WT and *Ckap2l^-/-^* P56 mouse brains stained with DAPI. Yellow dashed lines demarcate the boundaries of the cortices. Blue arrows point to the lobules that are underdeveloped in the cerebellum of *Ckap2l^-/-^*mice. (**H**) Western blot analysis of GLI3 and GLI3R levels in E14.5 WT and *Ckap2l^-/-^* mouse brains. Brains were dissected into dorsal and ventral region prior to lysis. (**I**) Quantification of GLI3R levels normalized to GAPDH. n = 4 biological replicates. Data are presented as mean ± SD. Statistics: two-tailed Student’s t-test. *, p < 0.05; ns, not significant.

**Figure S8.**
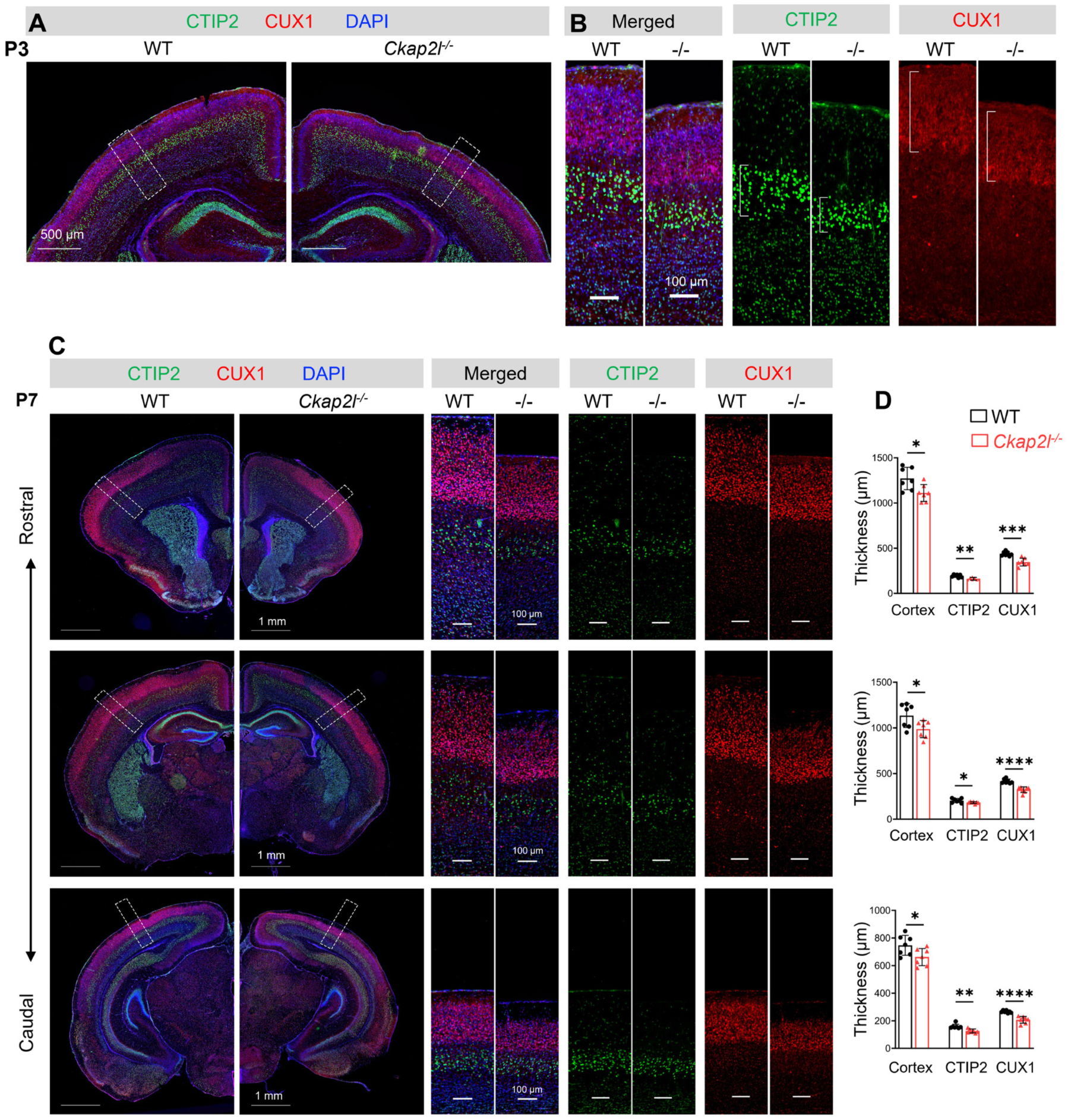
*Ckap2l^-/-^* mice exhibit reduced neuronal population. (**A**-**B**) Images of P3 brain sections stained with CTIP2, CUX1 and DAPI. Areas in the white dashed boxes are magnified and displayed in (**B**). (**C**) Images of P7 brain sections arranged from rostral to caudal positions. Sections were stained with CTIP2, CUX1 and DAPI. Areas in the white dashed boxes are magnified and displayed at the right side. (**D**) Quantification of the thickness of indicated layers in the corresponding rostral-caudal positions in (**C**). n= 7 brains were quantified for each genotype. Data are presented as mean ± SD. Statistics: two-tailed Student’s t-test. *, p < 0.05; **, p < 0.01; ***, p < 0.001; ****, p < 0.0001.

**Figure S9.**
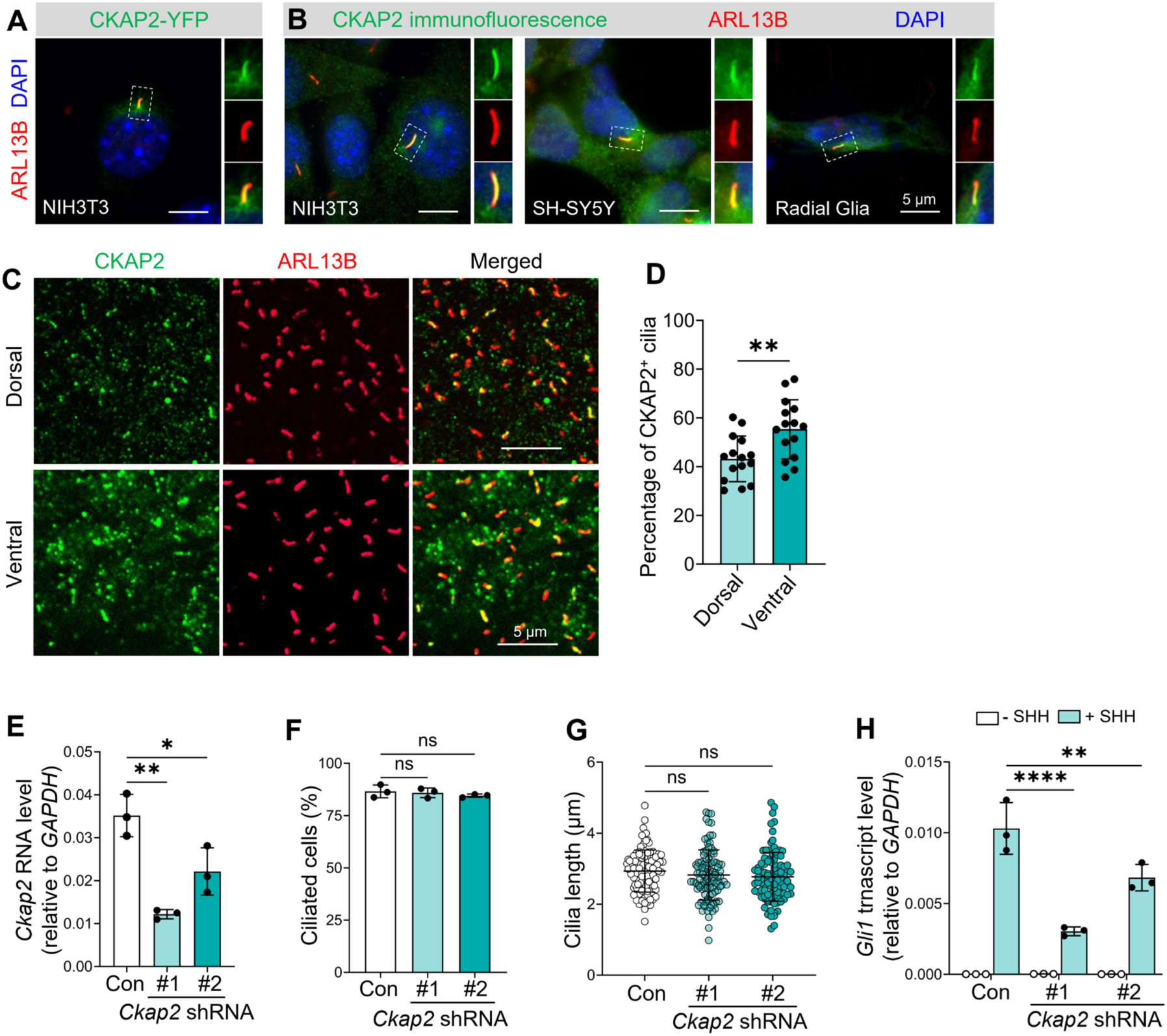
CKAP2 is present in the primary cilia and its loss impairs Hh signaling. (**A**) CKAP2-YFP is localized to the primary cilium in NIH3T3 cells. (**B**) Immunostaining with anti-CKAP2 antibody shows that endogenous CKAP2 is present in the cilia in multiple cell types, including NIH3T3 cells, SH-SY5Y and primary cultured radial glial cells. The cilium in the white dashed box is magnified and displayed at the right side. (**C**) *En face* views of whole-mount embryonic mouse brain following immunofluorescence staining with an anti-CKAP2 antibody. Endogenous CKAP2 is present in the primary cilia of E12.5 brains in both dorsal and ventral regions. (**D**) Quantification of CKAP2-positive cilia in (**C**). A total of 15 areas from 4 brains were quantified for the dorsal or ventral region. Data are presented as mean ± SD. Statistics: two-tailed Student’s t-test. **, p < 0.01. (**E**) *Ckap2* shRNA significantly reduced *Ckap2* gene expression. *Ckap2* mRNA levels were measured by qPCR. Control (con) shRNA is a non-targeting scrambled shRNA. Data are presented as mean ± SD. Statistics: One-way ANOVA with multiple comparisons (Tukey test). **, p < 0.01; *, p < 0.05. (**F**) Quantification of ciliated cells in control and *Ckap2* knockdown NIH3T3 cells. At least 100 cilia were quantified for each experimental condition. n=3 biological replicates. Data are presented as mean ± SD. Statistics: One-way ANOVA with multiple comparisons (Tukey test). ns, not significant. (**G**) Quantification of ciliary length in control and *Ckap2* knockdown NIH3T3 cells. A least 100 cilia were quantified for each experimental condition. Experiment was performed 3 times with similar results. Data are presented as mean ± SD. Statistics: One-way ANOVA with multiple comparisons (Tukey test). ns, not significant. (**H**) *Ckap2* knockdown attenuated SHH-induced Hh signaling. 3 days after lentivirus-mediated transfection of *Ckap2* shRNA, cells were serum-starved for 24 h with or without SHH. Hh signaling activity was evaluated via qPCR measuring transcript levels of *Gli1*. Data are shown as mean ± SD. Statistics: Two-way ANOVA with multiple comparisons (Tukey test). ****, p < 0.0001; **, p < 0.01.

